# Hypomorphic NOTCH1 Expression Alters Cardiomyocyte Cellular Architecture in Hypoplastic Left Heart Syndrome

**DOI:** 10.1101/2024.09.30.615809

**Authors:** Jordann Lewis, Travis B. Lear, Brent Schlegel, Dominic Woods, Krithika Rao, Amy Sentis, Jay Tan, Rajaganapathi Jagannathan, Zaineb Javed, Aine N. Boudreau, Timothy Nelson, Mousumi Moulik, Nadine Hempel, Bill B. Chen, Sruti Shiva, Dhivyaa Rajasundaram, Toren Finkel, Anita Saraf

**Affiliations:** Heart Institute, UPMC Children’s Hospital of Pittsburgh and UPMC Heart and Vascular Institute, Pittsburgh, PA 15312; Aging Institute, University of Pittsburgh/UPMC, Pittsburgh, PA 15219, USA; Department of Pediatrics, Children’s Hospital of Pittsburgh, University of Pittsburgh, Pittsburgh, PA 15312; Department of Medicine, University of Pittsburgh, Pittsburgh, PA 15213; Vascular Medicine Institute, Division of Cardiology, Department of Medicine, University of Pittsburgh, Pittsburgh, PA 15213; Department of Cell Biology, University of Pittsburgh, Pittsburgh, PA 15213, USA; Division of Pediatric Cardiology & Center for Regenerative Medicine, Mayo Clinic, Rochester, MN 55905; Department of Medicine, Division of Pulmonary, Allergy and Critical Care Medicine, Acute Lung Injury Center of Excellence, University of Pittsburgh, Pittsburgh, PA 15213, USA; UPMC Heart and Vascular Institute, Division of Cardiology, Department of Medicine, University of Pittsburgh, Pittsburgh, PA 15312

## Abstract

NOTCH1 is a protein involved in cardiac development and mutations in NOTCH1 are implicated in left sided congenital heart disease, including hypoplastic left heart syndrome (HLHS). Therapeutic advances for HLHS patients have been hampered by the absence of suitable experimental models. Here, using human induced pluripotent stem cells (hiPSCs), we have generated a series of CRISPR-edited complex heterozygous hypomorphic mutations in NOTCH1 that recapitulate mutations seen in HLHS patients. These cells demonstrate the downregulation of genes associated with mitochondria, the actin cytoskeleton, and cardiomyocyte development. In addition, these hypomorphic NOTCH1 hiPSCs cells showed an increased propensity to differentiate towards non-cardiomyocyte lineages such as fibroblasts and smooth muscle cells, a finding confirmed in HLHS patient-derived myocardial tissue. Abnormalities in sarcomeric and mitochondrial architecture contributed to decreased ATP production, abnormal calcium transients, and reduced contractility. Using a split nano-luciferase system as a NOTCH1 intracellular reporter, we screened a library of FDA-approved compounds. Auranofin, an agent currently employed for rheumatoid arthritis, was identified as a candidate drug that could rescue impaired differentiation and contractility seen in the hypomorphic NOTCH1 model. These findings point to cell-autonomous abnormalities in hypomorphic NOTCH1 pre-cardiomyocyte cells that can be leveraged to identify potential therapeutic strategies for patients with severe congenital cardiac abnormalities.

**Highlights:**

- CRISPR edited and patient-derived iPSCs with hypomorphic NOTCH1 expression were used to characterize a cell-based model of hypoplastic left heart syndrome.
- Hypomorphic NOTCH1 iPSCs have abnormalities in pathways associated with mitochondrial function, actin cytoskeleton, and cardiomyocyte development.
- Hypomorphic NOTCH1 iPSCs have less pluripotency and tend to skew differentiation away from cardiomyocytes and towards fibroblasts and smooth muscle cells.
- NOTCH1 is required for cardiac cytoskeletal and mitochondrial architecture, and to preserve contractility and ATP production.
- A high-throughput drug screen identified auranofin as an agent that may benefit patients with hypoplastic left heart syndrome.

## Introduction

Congenital heart disease (CHD) is the most common type of birth defect, affecting 40,000 newborn infants per year in the United States, with a total estimated prevalence of 2.4 million children and adults^1,2^. Complex congenital heart disease such as hypoplastic left heart syndrome (HLHS) is increasingly recognized as a product of compound heterozygous mutations that cumulatively lead to multiple structural defects at the vascular and myocardial level. While the severity of these abnormalities caused >90% mortality within the first month of life^3^, survival in these children has improved over the past five decades through iterative surgical operations^4^. However, while certain anatomic limitations have been addressed with surgeries, the underlying cardiomyopathy is pervasive and is the cause of >60% mortality in adulthood^5^. It has been challenging to study the pathophysiology of complex CHD like HLHS due to a lack of viable pre-clinical models and the difficulty acquiring human cardiac tissue. Induced pluripotent stem cells (iPSCs) offer a way to overcome these limitations and are increasingly being used to investigate phenotypic changes associated with specific gene mutations in CHD^6–12^. Specifically, use of patient-derived iPSCs as well as gene-edited iPSCs can provide a platform to study the link between known genetic mutations among CHD patients and correlate the resulting phenotype at a molecular and cellular level.

In this study, we genetically engineered compound heterozygous mutations in various domains of NOTCH1^13,14^, -a developmental protein implicated in HLHS, to understand their phenotypic correlation with cardiomyocyte (CM) physiology. NOTCH1 is a developmental protein involved in myocardial, ventricular, and outflow tract development and is necessary for valvulogenesis, proper trabeculation of the ventricular chambers, and formation of endocardial cushions/AV canal/septa^15–18^. NOTCH1, a transmembrane protein, is primarily activated during embryogenesis and has distinct signaling pathways activated through its extracellular (ECD) and intracellular (ICD) domains. While the ECD interacts with ligands from neighboring cells and is involved in juxtracrine signaling, the ICD domain is released after proteolytic cleavage and alters gene expression by acting as a transcription factor by binding to the DNA-bound RBP-JƘ protein complex^19^. Compound heterozygous single nucleotide mutations involving both ECD and ICD were observed and characterized in a family with an HLHS proband^20^, where the paternal allele had a mutation in the extracellular domain (ECD) (P1256L) and the maternal allele had a mutation in the intracellular domain (P1964L). In this family, the mother had a bicuspid pulmonary valve, while the father has no structural heart disease, and the proband has HLHS. Given its ubiquitous involvement in the development of the heart, it is not surprising that different mutations in NOTCH1 lead to varying degrees and severity of cardiac abnormalities^8,20,21^.

To further evaluate the role of hypomorphic NOTCH1 mutations in cardiomyocyte signaling and function in HLHS, we used Clustered Regularly Interspaced Short Palindromic Repeats (CRISPR)/CRISPR-associated protein 9 (Cas9) gene editing to create a wide range of isogenic clones of human iPSCs with a wide range of NOTCH1 hypomorphic expression through mutations in the intracellular and extracellular domains. Replicating suspected pathogenic mutations in hiPSCs with an independent genetic background can isolate the effect of the mutations from the genetic background of the proband. Hence, to further understand the pathogenicity of the hypomorphic NOTCH1 mutations, we compared the phenotype with iPSCs derived from the proband described above^20^. Our studies show distinct phenotypic correlations with ECD and ICD NOTCH1 complex heterozygous mutations in 1) downstream gene regulation patterns and differentiation and 2) differential regulation of mitochondrial and cytoskeletal proteins. Phenotype-genotype correlation is also seen after differentiation whereby 1) lineage mapping shows preferential differentiation of hypomorphic NOTCH1 into the fibroblasts (FB) and smooth muscle cells (SMC) and 2) cytoskeletal abnormalities in cardiomyocytes which cause a significant overlap of CM, SMC, and FB. These results suggest that hypomorphic NOTCH1 expression leads to abnormalities at the stem/progenitor state that influence the morphology and function after differentiation and affect the proband through a similar mechanism. Finally, we screened a library of FDA-approved compounds using a high-throughput quantitative split-Nano luciferase system to identify agents that augment NOTCH1 signaling. Using a combination of optical and fluorescence-based readouts, we identified auranofin as an agent that can rescue the functional and structural defects seen in our hypomorphic NOTCH1 iPSC-CMs.

## Results

### Heterozygous NOTCH1 mutations cause hypomorphic expression of NOTCH1 and downstream proteins

CRISPR/Cas9 gene editing of NOTCH1 was used to generate multiple independent iPSC cell types with compound heterozygous genetic patterns of hypomorphic NOTCH1 expression (Figure 1A). Based on previous observations, we sought to target both the extracellular domain (ECD) and intracellular domain (ICD) of NOTCH1^20^. While the mutations in the patient trio (parents and proband) were single nucleotide polymorphisms (SNPs), these mutations did not independently show significant cardiac abnormalities. Hence, while we targeted the same regions of NOTCH1 with CRISPR editing, we sought to create more elaborate alterations to understand the relative implications of mutations in each protein domain. Multiple NOTCH1 hypomorphic clones (HC) were sequenced, and clones with homozygous knockout mutations within the targeted NOTCH1 domain were excluded. Two clones with compound heterozygous mutations within the NOTCH1 extracellular (ECD) and intracellular (ICD) domain each were selected for further evaluation (Figure 1B). Markers of pluripotency KLF4, OCT3/4, SOX2, and MYC were downregulated in NOTCH1 iPSC hypomorphic clones. However, there were differences in the relative degree of downregulation between the clones (Figure 1C). Location and severity of the NOTCH1 mutation influenced NOTCH1 expression and, as expected, decreased expression of NOTCH1 target genes, including Hey1, Hes1, and Hey L (Figure 1D). Notably, downstream expression of these NOTCH1-regulated genes was differentially affected amongst the various hypomorphic clones. NOTCH1 hypomorphic clones showed decreased differentiation into CMs as compared to WT iPSCs (Figure 1E). Clones with heterozygous mutations leading to intermediate NOTCH1 expression showed reduced differentiation efficiency as determined by NKX2.5 expression, whereas hypomorphic clones with stop codons or frame-shift mutations in both alleles resulted in a complete loss of viability following initiation of differentiation.

**Figure 1:**
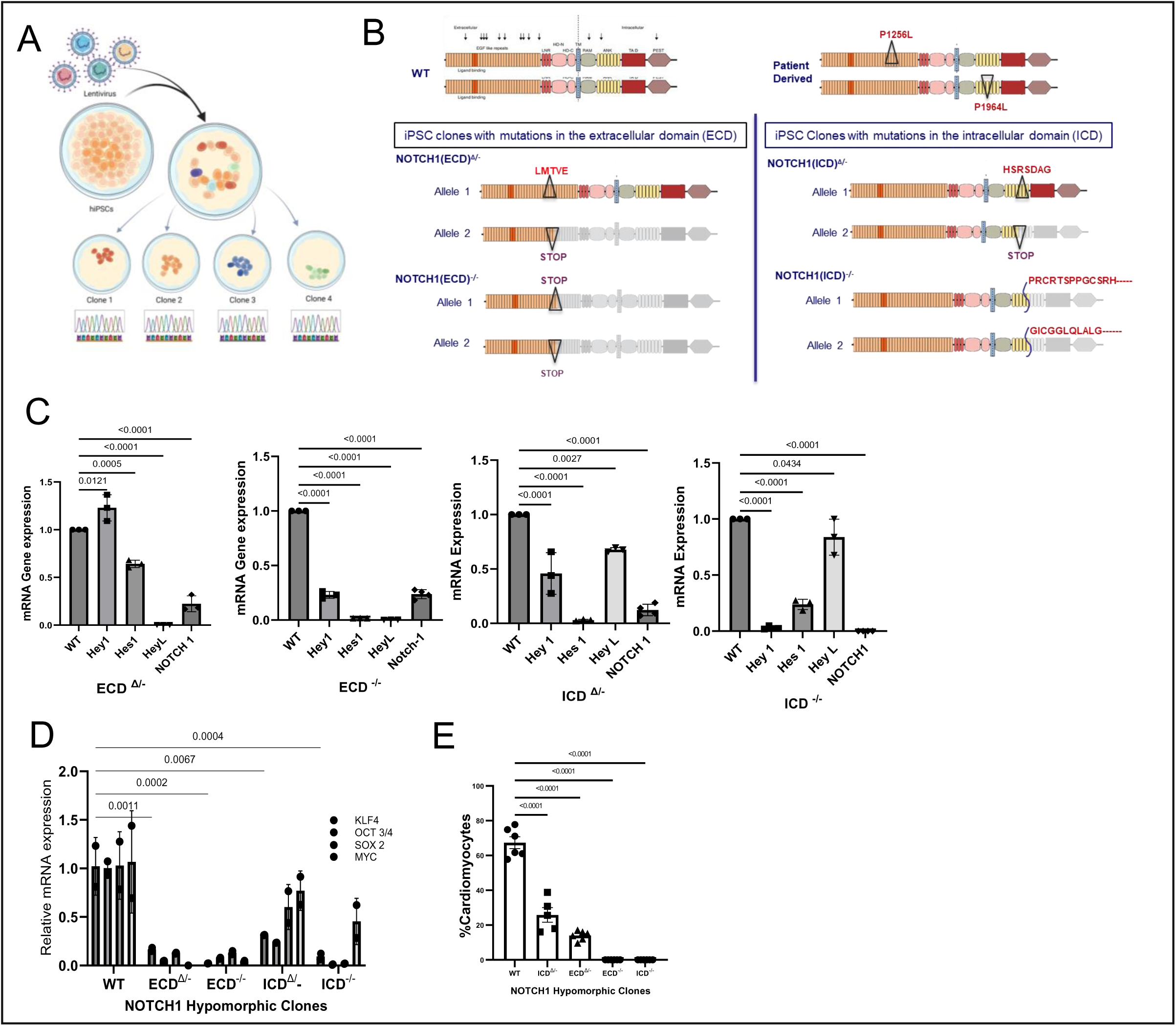
Creation and characterization of hypomorphic NOTCH1 hiPSC. (A) Schematic for generating hypomorphic hiPSC clones. hiPSCs were edited using CRISPR technology using guide RNAs for the NOTCH ECD and ICD domains. After selection, individual cells were picked and expanded independently into mutated clonal populations after which they were sequenced (B) Schematic and nomenclature of NOTCH1 mutations in hiPCs clones. DNA sequencing characterized two colonies from each mutation product in the ICD and ECD domains. Arrows associated with WT cells represent the location of mutations noted in NOTCH1 in other congenital heart disease patients. HLHS patient-derived iPSC cell line carried a complex heterozygous mutation in ICD and ECD domains of NOTCH1. (C) Pluripotency markers in hypomorphic NOTCH1 hiPSCs were significantly reduced compared to WT iPSCs. (n=3 clones per mutation product) (D) Downstream NOTCH1-regulated proteins, including Hey1, Hes1, and HeyL, showed a significant decrease in expression in hypomorphic NOTCH1 iPSCs as compared to WT iPSCs. (n=3-4 per each clonal group). (E) Differentiation of hiPSCs with and without hypomorphic NOTCH1 expression showed that hypomorphic NOTCH1 expression decreased the percentage of cells that differentiated into cardiomyocytes. Cells with premature stop codons in both alleles of ECD and ICD could not differentiate into cardiomyocytes, although their corresponding hiPSCs were viable. (n=4 clones per mutation product with multiple regions of interest per well).

### Hypomorphic NOTCH1 leads to cytoskeletal dysregulation and mitochondrial stress

NOTCH1 is a complex and large protein with multiple signaling pathways that are regulated by its various domains. Previous studies in NOTCH1 deficient CMs have demonstrated a higher prevalence of heterogenous myofilament organization^20^. Bulk RNAseq of CRISPR engineered ECD^Δ/-^, ICD^Δ/-^, and WT hiPSCs were performed to elucidate pathways that are differentially regulated within ECD^Δ/-^ and ICD^Δ/-^ iPSC clones (Figure 2). The principal component analysis demonstrated that ECD^Δ/-,^ ICD^Δ/-^, and WT were distinguishable from each other, with the most disparate distribution among the ECD^Δ/-^ and WT iPSC clones (Supplemental Figure 1A). Volcano plot analysis demonstrated a distinct set of genes being differentially expressed in the ECD^Δ/-^ and ICD^Δ/-^ clones, as compared to WT hiPSC. However, Microtubule Associated Protein 2, (MAP2) a cytoskeletal protein that connects microtubules and intermediate filaments^22^ was the most significantly and equally downregulated in both hypomorphic clones. Additionally, Sox1, an ectoderm marker in fetal development, was equally and strongly downregulated in both hypomorphic clones. In addition, a unique set of genes was found to be differentially regulated in each hypomorphic clones as shown in the volcano plot (Figure 2A). As represented in the Venn diagram (figure 2B), 11791 genes were differentially regulated as compared to WT hiPSCs encompassing various pathways associated with metabolic and structural proteins. A subset of 219 genes were specifically regulated in the ECD^Δ/-^, whereas 158 genes were specifically regulated in ICD^Δ/-^. Analysis of specific pathways differentially regulated between ECD^Δ/-^ and ICD^Δ/-^ identified molecules associated with CM cytoskeletal signaling (Actin, RAC), mitochondrial stress (NRF2, eIF2, renin-angiotensin), bioenergetics (HIF1α, AMPK, mTOR), cell-cell signaling (integrin, adherens junction, paxillin) and cardiac differentiation (WNT/β-catenin, PDGF, mTOR, PAK, VEGF, and TGFβ) (Figure 2C). Mitochondrial pathways associated with oxidative stress and bioenergetics (eIF2, NRF2, renin-angiotensin, HIF1α) were more robustly dysregulated in ECD^Δ/-^ hypomorphic clones, whereas ICD^Δ/-^ hypomorphic clones showed a slightly higher propensity to increased dysregulation associated with cytoskeletal protein and their related cell-processes including migration, proliferation, differentiation (Integrin, paxillin, ERK/MAPK, Wnt/β- catenin) and cell interaction and mechano-transduction (integrin and paxillin). Of the top 20 genes associated with each of these pathways (Figure 2D), most genes that were similarly regulated in both hypomorphic clones as compared to WT (ACTC1, TRIOBP, ANK2, BIABP2L1, MYO1C, LMNA, RAC3) were those associated with cytoskeletal signaling and organization.

**Figure 2:**
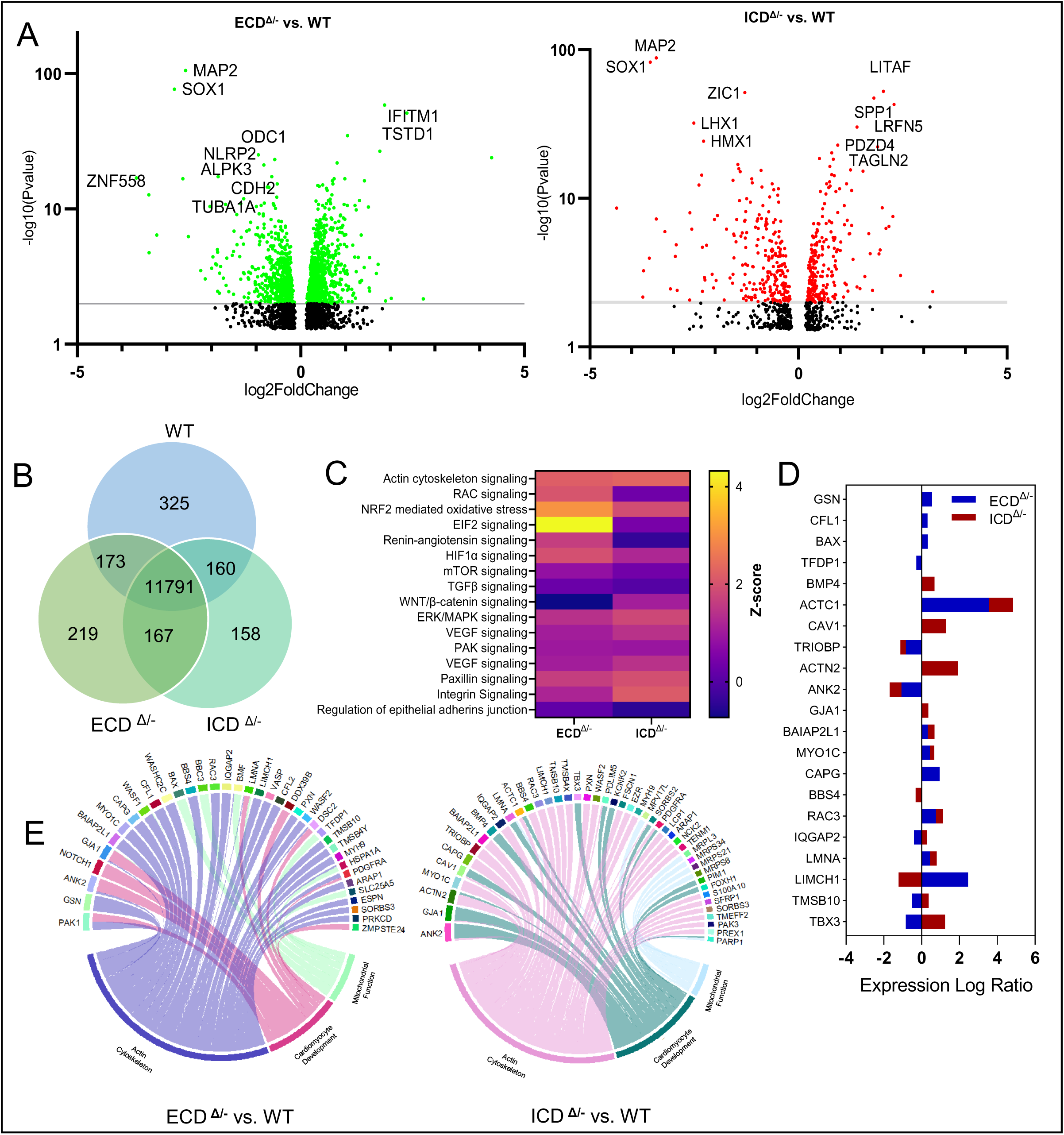
Transcriptomics of WT and hypomorphic NOTCH1 hiPSC: (A) Volcano plots showing differential gene expression of ECD Δ/- vs. WT and ICD Δ/- vs. WT. (n=3 clones per mutation product). (B) Venn diagram for differentially expressed genes between WT, ECD^Δ/-^ and. ICD^Δ/-^ (C) Differentially affected signaling pathways between hypomorphic NOTCH1 iPSC clones with ECD^Δ/-^ vs. ICD^Δ/-.^ (D) Profile of top 20 transcriptomic markers differentially regulated between ECD^Δ/-^ vs. ICD^Δ/-^ in clones of hiPSCs (E) Cord diagrams that show differentially expressed genes associated with mitochondrial function, actin cytoskeleton and cardiomyocyte development in ECD^Δ/-^ vs. WT and ICD^Δ/-^ vs. WT in iPSCs.

Genes associated with cell stress, autophagy, and apoptosis (BAX, IQGAP2, LMNA) were more robustly associated with ICD^Δ/-.^ To evaluate the relative contribution of dysregulated gene expression with the respective pathways involved in cardiomyocyte development, chord diagrams were generated for the top-20 regulated genes in each of the two hypomorphic clones (Figure 2E). Both ECD^Δ/-^ and ICD^Δ/-^ clones showed a significant dysregulation in pathways associated with actin-cytoskeleton formation, mitochondrial function, and cardiomyocyte development. GO enrichment analysis scatter plot of ECD^Δ/-^ vs. WT and ICD^Δ/-^vs WT showed similar dysregulation of protein synthesis pathways for both mutations within ECD and ICD loci as compared to WT. However, ECD^Δ/-^ showed significant differences in oxidoreductive activity and calcium signaling in addition to multiple developmental pathways (Supplemental Figure 1B).

### Unselected differentiation of hypomorphic NOTCH1 iPSCs gives rise to heterogeneous cardiac cells

Given the differences in signaling pathways noted in ICD versus ECD hypomorphic NOTCH1 clones, we next sought to differentiate these cells into CMs. iPSCs differentiate into a heterogenous cell population in the absence of glucose restriction or lactate enrichment, strategies commonly incorporated into CM differentiation protocols^23^. To investigate the cellular heterogeneity amongst post-differentiated cells, we performed single-cell transcriptomics on unselected products of differentiation and analyzed them at day 25-27 post-differentiation using 10X genomics (Figure 3A). Manifold Approximation and Projection (UMAP) algorithm revealed 7 cell clusters, including cardiomyocytes (CM), fibroblasts (FB), smooth muscle cells (SMC), non-specific proliferating cells (NPC), fibroblast-like proliferating cells (FPC), endothelial (EDC) and epithelial cells (EPC) (Figure 3B). Top 20 gene markers for each cell cluster are noted in supplemental table 2 and were used to determine the phenotypes of cells within each cluster. Representative genetic signatures of aggregate cells are visualized in the heat map (Supplemental Figure 2A), which shows that while most markers within each cluster were specific to the cell type, some markers were shared across multiple clusters. This was more apparent with endothelial cells, fibroblast-like proliferating cells, fibroblasts, non-specific proliferating cells, and smooth muscle cells. The presence of proliferating cells and cells with ambiguous phenotypes are common in unselected iPSC differentiation products^24^. Violin plots of representative cells with their markers are noted in Figure 3C. Proliferating cells (PC, i.e., FPC and NPC) cells shared genetic markers with multiple clusters but more significantly expressed markers of proliferation (e.g., TPX2, HMGB2, CDK1, and ORC6). As compared to non-specific PC, FPC showed more robust expression of markers that were representative of invasive proliferating fibroblasts (SRFRP5, TM4SF1, SPRR2F). CM markers represented both atrial and ventricular cells (atrial: MYL7, MYH6; ventricular: TNNT1, TNNT2, MYH7, and TNNC1) but also expressed markers that overlapped with SMC (ACTA2, ACTN1) and fibroblasts (DCN and COL1A1, PDGFRA) (Figure 3D). Differentiated cells derived from NOTCH1 hypomorphic clones showed a significant change in phenotype from WT cells in two major categories. Secondly, within the CM subgroup, while cells were found to have CM markers (TNNT1, MYL4, TNNT2, TNN, and ACTC1), ECD^Δ/-^ and ICD^Δ/-^ hypomorphic clones CMs had markers that overlapped with SMCs (ACTA2, ACTN1, DCN, COL1A1, PDGFRA and SFRP5) and FBs (DCN, COL1A1, PDGFRA) suggesting that hypomorphic NOTCH1 CMs also have FB and SMC characteristics, consistent with previous analysis^7,20,25^. While WT CMs had a lower expression of SMC and FB genes (PDGFRA, DCN, FN1), these markers were significantly increased in ECD^Δ/-^ and ICD^Δ/-^ HC (Figure 3D, Supplemental Figure 2B). We performed gene set-enrichment analysis (GSEA) per cell type for each cell cluster (Figure 3E). GESA demonstrated that genes in pathways associated with sarcomere organization and actin cytoskeleton were preferentially downregulated in ECD^Δ/-^ CM, SMC, and EDC. Non-CM cells (FB and NSP) showed decreased expression of sarcomere organization and actin cytoskeleton in both ECD^Δ/-^ and ICD^Δ/-^ hypomorphic clones, indicating that abnormal cytoskeletal architecture of CM-associated myocardial cells may also contribute to disease pathology. CM associated with ECD^Δ/-^ showed a greater downregulation in multiple additional pathways (such as adrenergic signaling, oxidative phosphorylation, cardiac muscle contraction, and cardiac tissue morphogenesis) as compared to WT and ICD^Δ/-^ and an upregulation in catabolic process and insulin-like growth factor receptor (IGFR) signaling, which may contribute to abnormal energy production and utilization within hypomorphic NOTCH1 CMs. NSP, representing progenitor cells that give rise to differentiated myocardial cells in both ECD^Δ/-^ and ICD^Δ/-^ HC, showed a significant increase in canonical and non-canonical Wnt signaling, which is implicated in the development of the anterior or 2^nd^ heart field, consistent with the idea that second heart field development is significantly enhanced due to downregulation of the development of the NOTCH1 dependent 1^st^ heart field^18^ (Figure 3E). Overall, our sc-transcriptomics data indicates that hypomorphic NOTCH1 expression in differentiating cardiac cells affects the contractile and cytoskeletal.

**Figure 3:**
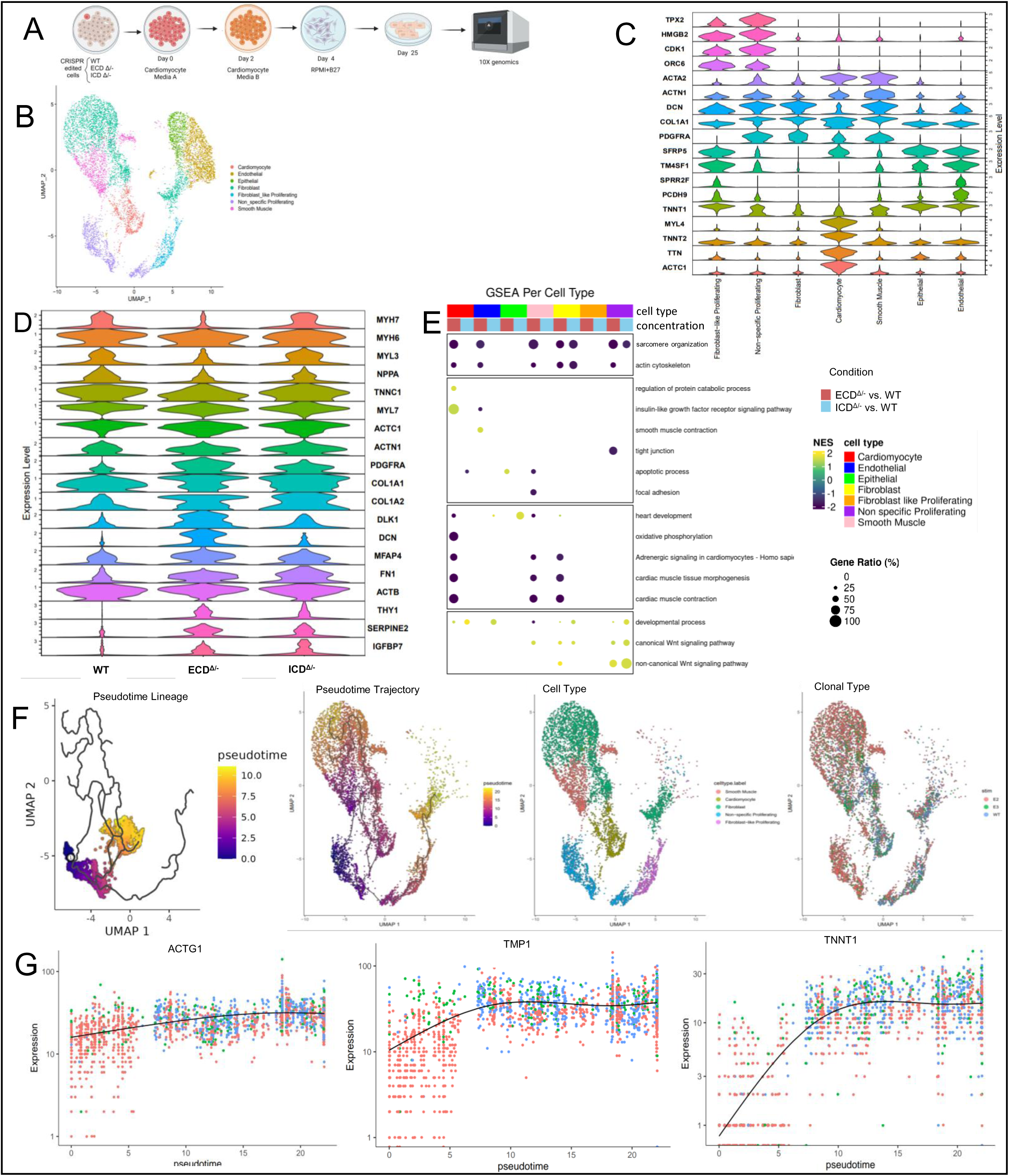
Single-cell transcriptomics demonstrated that hypomorphic NOTCH1 iPSCs differentiated preferentially into non-cardiomyocyte cell types. (A) schematic for differentiation of iPSCs into myocardial organoids for sc-transcriptomics analysis. Cells were analyzed with 10x genomics on Day 25 after initiation of differentiation s (B) UMAP of cells resulting from the differentiation process showed 7 clusters of cells including fibroblasts (FB), smooth muscle cells (SMC), epithelial (EPC) and endothelial cells (EDC) (n=3 replicates from each clone per condition). (C) Violin plots of cell-specific molecular markers of cell subsets identified by UMAP of sc-transcriptomics (D) Violin plots of cell-specific molecular markers of CM subset between ECD^Δ/-^, ICD^Δ/-^ and WT cells show an increase in SMC and FB markers in ECD^Δ/-^, ICD ^Δ/-^ as compared to WT cells. (E) GSEA analysis shows significant abnormalities in pathways associated with sarcomere assembly and cardiomyocyte development in ECD^Δ/-^ as compared to WT. ICD^Δ/-^ CMs showed differences in pathways associated with apoptosis in CMs, and in pathways associated with non-cardiomyocyte cells including FB and PNS cells. (F) UMAP projections showing all calculated Monocle pseudotime trajectories of differentiating cells, originating from the Non-specific Proliferating cluster and extending towards smooth muscle, cardiomyocyte, and fibroblast clusters. Darker colors indicate less differentiation (similarity to the non-specific cluster), while lighter colors indicate further distance along the pseudotime trajectory (higher differentiation). UMAP projections colored by clone type and annotated cluster are also shown for additional comparison. (G) Pseudotemporal dynamics of a selected lineage extending from the Non- specific proliferating cluster to the cardiomyocyte cluster. Scatter plots show gene expression for key developmental markers as a function of pseudotime, highlighting increased expression of differentiation markers ACTG1, TMP1, TNNT1 and increased representation of NOTCH1 hypomorphs further along the calculated trajectory. Points are colored by their respective clone type. The line indicates the mean trend of expression along the pseudotime lineage.

Next, we performed pseudotime trajectory analysis to reveal the progression of differentiation from progenitor cells to cardiomyocytes during iPSC differentiation and identified four different lineage pathways (Supplemental Figure 3). This analysis revealed a higher incidence of cardiomyocytes differentiating along the Wnt signaling pathway in ECD^Δ/-^ and ICD^Δ/-^ HC WT iPSCs as compared to WT HC. Similarly, markers associated with actin organization in CMs including γ-cytoplasmatic actin (ACTG1), Tropomyosin I (TMP1) and Troponin T Type 1 (TNNT1) increased in WT cells during the later stages of differentiation in the psuedotime trajectory (Figure 3G). However, similar increases were not seen with ECD^Δ/-^ and ICD^Δ/-^ HC. (Supplemental Figure 3B, 3C)

### Increased mitochondrial fission and decreased ATP synthesis in hypomorphic NOTCH1 CMs

Previous studies with HLHS patient-derived iPSCs demonstrated abnormalities in mitochondrial size and inner membrane structure by electron microscopy^7^. We therefore sought to investigate mitochondrial morphology in our models. CMs derived from NOTCH1 hypomorphic clones showed abnormal mitochondrial morphology and function (Figure 4). Mitotracker fluorescence analysis of ECD^Δ/-^ and ICD^Δ/-^ CMs demonstrated an increased pattern of fission as compared to WT CMs (Figure 4A). Multiple mitochondrial characteristics were analyzed (Figure 4B); WT CMs showed an increase in the number of mitochondrial branches/mitochondria as compared to ECD ^Δ/-^and ICD^Δ/-^ (p=0.0002). Overall, the mitochondrial footprint was greater in WT CM as represented by the mitochondrial perimeter (5.261 µm in WT CM) vs. hypomorphic clones (3.469 with ECD^Δ/-^ vs. 3.063 with ICD^Δ/-^ p=0.0003). Additional mitochondrial parameters, including mitochondrial area/count (p=0.0003), total branch length (p=0.0002), and form factor, which represents complexity and branching of mitochondria (p=0.0002), indicated shorter, less branched, and smaller surface area covered by mitochondria in CMs derived from the hypomorphic clones. Mitochondrial abnormalities were evident in NOTCH1 iPSCs before their differentiation as noted through the hiPSC transcriptomics data. ECD^Δ/-^ CMs showed a significant increase in ROS levels as compared to WT CM (p<0.0001). However, increased oxidative stress was not seen in ICD^Δ/-^ iPSC-CMs (p=0.075) compared to WT (Figure 4C and 4D). Mitochondrial Seahorse analysis showed a significant decrease in oxygen consumption rate (OCR) in ICD^Δ/-^ > ECD^Δ/-^ > WT cells (Figure 4E). However, mitochondrial content between cells did not show a significant difference between WT and NOTCH1 hypomorphic CMs (Figure 4F). Hence, hypomorphic NOTCH1 expression in CMs showed increased fission and decreased ATP production. Consistent with our transcriptomics analysis, CMs from ECD^Δ/-^ iPSCs showed an increase in ROS expression suggestive of abnormal mitochondrial development and function post differentiation.

**Figure 4:**
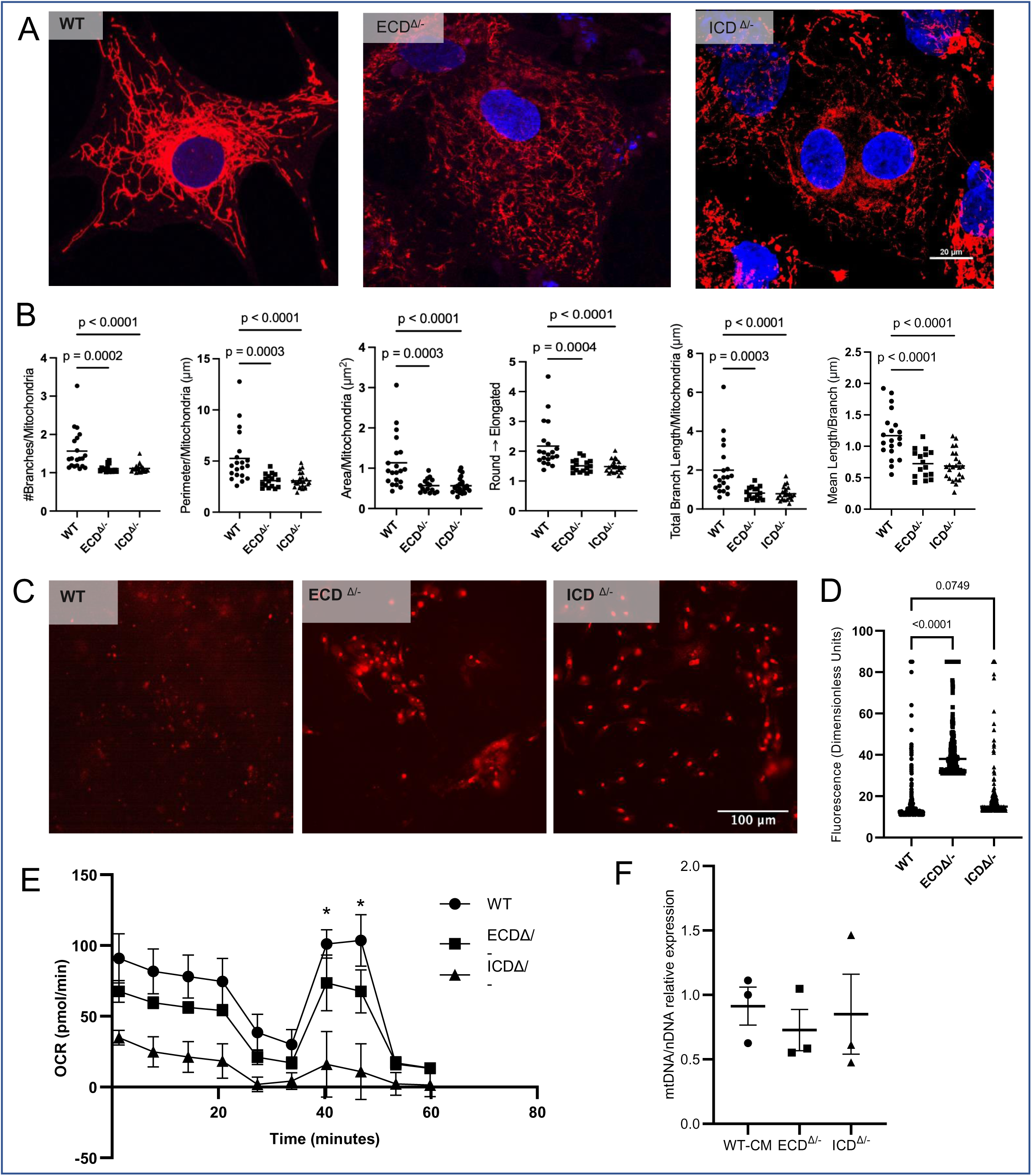
Mitochondrial structure and function is abnormal in hypomorphic NOTCH1 iPSC-CMs. (A) Mitochondrial architecture in WT iPSC-CMs shows an extensive intracellular reticular pattern that is disrupted in hypomorphic iPSC-CMs. (B) Quantitative analysis of iPSC-CMs show decreases in mitochondrial branches/mitochondria, mitochondrial perimeter count, mitochondrial area/count, mean form factor (where higher values represent more linear mitochondria) and total branch length (n=3-4 clones per mutation product). (C,D) Reactive oxygen species production increases significantly in hypomorphic NOTCH1 iPSC-CM clones as compared to WT alone (n=3 clones per mutation product). (E) Sea-horse assay shows a decrease in ATP production in hypomorphic NOTCH1 iPSC-CMs as compared to WT (n=6 clones per mutation product per time point). (F) The overall mtDNA expression does not significantly change between WT and ECD^Δ/-^ and ICD ^Δ/-^ CM (n=3 clones per mutation product).

### Sarcomeric architecture in hypomorphic NOTCH1 cardiomyocytes shows increased disorganization and absence of z-discs

Histological sections of myocardial tissue from HLHS patients show disorganized sarcomeres and an aberrant organization of cardiomyocytes which leads to abnormal calcium transients and contractility^26^. Numerous studies of iPSC-derived CMs from HLHS patients show a similar pattern of disorganization. Similarly, NOTCH1 CMs showed significant abnormalities in sarcomeric architecture, calcium transients, and contractility^7,20^. Fluorescence staining of α-actinin in 25-day post-differentiation hiPSC-CMs showed an organized compact sarcomeric architecture in WT cells (Figure 5A). However, hypomorphic NOTCH1 iPSCs-CMs showed a variety of abnormalities including loss of striations (Z-bands) in significant sections within the sarcomeres of the cardiomyocytes, decreased packing density, disorganization of sarcomeres and presence of non-filamentous α-actinin (Figure 5A, 5C). The average sarcomere organization score for WT CM was 2.51, as compared to the NOTCH1 hypomorphic clones which were ECD^Δ/-^ 1.2 and ICD^Δ/-^ 1.25. These observations indicate that hypomorphic NOTCH1 hiPSC-CMs exhibit a significant decrease in the organization, formation of filaments, and z-discs of functional sarcomeres (Figure 5B). Live cell imaging of fluorescent alpha-actinin-tagged sarcomeres showed WT CMs had filamentous sarcomeres with z-discs, extending from the nucleus to the cell boundary. The z-discs are organized parallel to each other and beat synchronously. In ECD^Δ/-^ and ICD^Δ/-^ CM, live-cell imaging showed that organization, as well as z-disc architecture, was disrupted, with long, non-striated filaments spanning the cells. As a result, only discrete sections of the cells with striated sarcomeres exhibited contractility, whereas the section with non-striated sarcomeres did not contract (Figure 5C, Movies 1,2,3).

**Figure 5:**
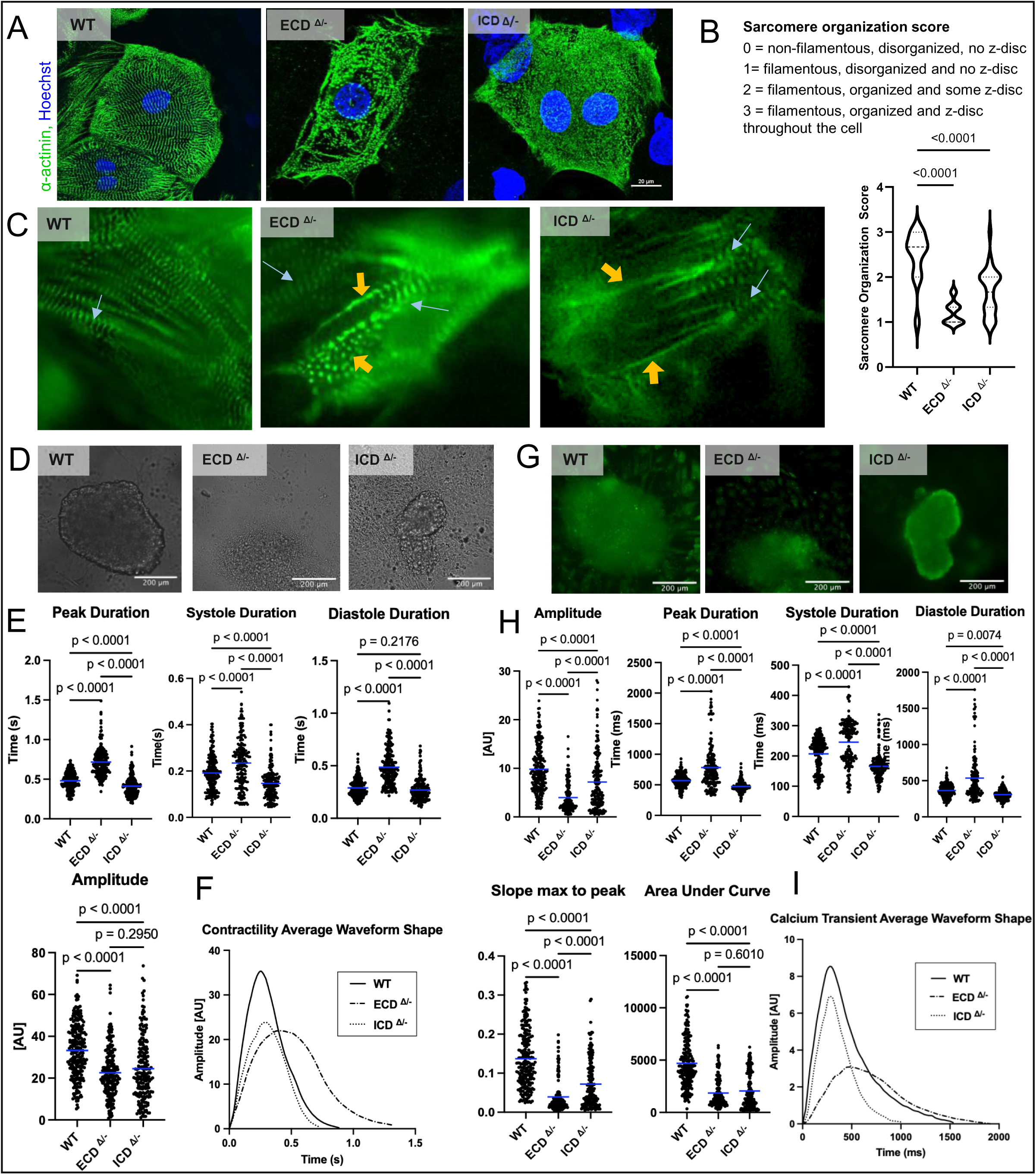
Sarcomeric architecture is disrupted in HM NOTCH1 mutations. (A) Representative CMs from WT, ECD^Δ/-^ and ICD^Δ/-^ demonstrate an increasingly filamentous and disorganized sarcomeric architecture in NKX2.5+ cells. (B) Sarcomere organization score showed a lower score for both ECD^Δ/-^ and ICD^Δ/-^ CMs. (n=50 cardiomyocytes from each clone per condition) (C) Live-cell imaging of GFP-tagged α-actinin demonstrated loss of striations and abnormal contractility of ECD^Δ/-^ and ICD^Δ/-^ CMs as compared to WT CMs. (D) Bright-field imaging of organoids formed from WT, ECD^Δ/-^ and ICD^Δ/-^ were analyzed for contractility and showed a decrease in amplitude between WT and ECD ^Δ/-^ CMs. (E) A significant increase in peak duration, and duration of systole and diastole with ECD^Δ/-^ and a decrease with ICD^Δ/-^ as compared to WT was noted. (n= 209-245 cells from 7-10 organoids from 3 clones for each mutation). (F) Representative beating waveforms of WT, ECD^Δ/-^ and ICD^Δ/-^ represented from quantitative data obtained from cardiac organoids of WT and hypomorphic NOTCH1 clones. (G) Corresponding representative Fluo-4 calcium transients of beating organoids represented in (D). (H) The amplitude of calcium flux of both ECD^Δ/-^and ICD^Δ/-^ organoids showed a significant decrease as compared to WT. Peak duration and duration of systole and diastole of ECD^Δ/-^ was increased as compared to WT organoids and decreased in ICD^Δ/-^ as compared to WT. (n= 170-238 cells from 7-10 organoids from 3 clones per mutation product) (I) Representative fluorescent waveforms of WT, ECD^Δ/-^ and ICD^Δ/-^.

### Hypomorphic NOTCH1 organoids showed abnormal patterns of beating and calcium dynamics

Patients with HLHS have an increased predisposition to arrhythmia and sudden cardiac death, which suggests that these patients have an inherently abnormal electromechanical coupling in the heart^27^. Krane et al demonstrated multiple abnormalities in calcium handling in hiPSCs derived from patients with HLHS^28^. In our studies, organoids assembled from ECD^Δ/-^ and ICD^Δ/-^ CM showed significantly different patterns of beating as compared to WT organoids (Figure 5D). Duration of systole in ECD^Δ/-^ CM was significantly longer than WT CM (0.236s vs 0.192s) (p<0.0001). Similarly, the duration of diastole in ECD^Δ/-^ CM was also longer (0.487s) as compared to WT CM (0.487s vs. 0289s, p<0.0001), as was total peak time (ECD ^Δ/-^ CM = 0.718s vs. WT CM = 0.481s, p<0.0001) indicating prolonged contraction and relaxation, and reduced frequency. On the other hand, ICD^Δ/-^ organoids had the lowest durations for systole and total peak time, indicating a beating frequency greater than WT (Figure 5E). However, these faster beats may have been less effective, as indicated by their inability to achieve as high a systolic amplitude as WT CMs. In fact, both ICD^Δ/-^ and ECD^Δ/-^ organoids exhibited significantly reduced amplitudes as compared to WT organoids (p<0.0001) (Figure 5F). Calcium transients were also abnormal in hypomorphic NOTCH1 organoids, following a similar pattern as seen with contractility. ECD^Δ/-^ organoids had the slowest, and ICD^Δ/-^ organoids the fastest calcium dynamics respectively (Figure 5H). Calcium efflux corresponding to the duration of systole was longest with ECD^Δ/-^ CM vs. WT CM (245.5ms vs. 165.7ms) and longer than that of ICD^Δ/-^ CM (165.7ms), (p<0.0001). Calcium influx corresponding to duration of diastole was also the longest for ECD^Δ/-^ CM as compared to WT CM (536.2 ms vs 363.7ms (p<0.0001)). ECD^Δ/-^ hypomorphic clones also had the longest total duration of calcium flux ECD^Δ/-^ CM as compared to WT CM (781.7ms vs. 570.7 ms), (p<0.0001). ECD^Δ/-^ and ICD^Δ/-^ organoids both had significantly reduced amplitudes as compared to WT CM (p<0.0001) and maximum slope to peaks (p<0.0001). Areas under the curve, which represents total calcium flux/signal/magnitude were also greatest for WT CM as compared to NOCH1 hypomorphic clones (p<0.0001) (Figure 5I).

### Differentiated iPSCs from HLHS patient phenocopy CRISPR-edited NOTCH1 hypomorphic cells

Colonies of iPSCs from the parents as well as the affected HLHS proband were previously genetically and functionally characterized with respect to cellular physiology and electromechanical properties^20^. These studies showed that as compared to parental iPSCs, proband hiPSCs showed decreased expression of NOTCH1 and downstream genes Hes1 and Hey1, impaired pluripotent stem cell markers, and decreased cardiogenesis when compared to the non-HLHS parental cells^20^. Additionally, proband hiPSC-CMs also demonstrated an increase in disorganized myofibrils^20^. We could recapitulate these observations in our studies (Figure 6). All iPSC CMs clones from parents and probands were able to form iPSC colonies (Figure 6A). Immunofluorescence (IF) staining of CMs derived from parental hiPSCs showed a well-organized dense sarcomeric pattern of distribution, whereas CMs from NOTCH1^Δ^/ ^Δ^ proband derived cells showed decreased sarcomeric density (Figure 6B). Transcriptomics analysis of proband iPSCs showed dysregulated GO pathways overlapping with CRISPR-edited iPSCs (Figure 6C). Similar to ECD^Δ/-^ iPSCs, the probands iPSCs showed a significant dysregulation with eIF2 signaling, NRF2 mediated oxidative stress, actin cytoskeletal signaling, WNT/β-catenin signaling, and HIF1α signaling. Similar to ICD^Δ/-^ iPSCs, the probands iPSCs showed dysregulated RAC and renin-angiotensin signaling. There was notable dysregulation of epithelial adherens junctions noted with proband iPSCs, that were not as dysregulated with ECD^Δ/-^ or ICD^Δ/-^ iPSCs. Differentiated unselected iPSCs showed a similarly diverse population of cells including CMs, EDCs, EPCs, SMCs, FB and FPCs, and NSPs. Similar to our CRISPR-edited cells, GSEA Per Cell Type analysis showed multiple pathways associated with actin cytoskeletal and sarcomere organization were altered within proband iPSC-CMs (Figure 6D). CMs had dysregulated cardiac function and heart processes and regulation of ATP-dependent activity. Non-CMs showed significant abnormalities across various pathways including cytoskeletal organization, formation of cardiac structures, and development and increase in oxidative stress-related pathways, predominantly in SMC and FB cells, but also across EPCs. Lastly, we compared our single-cell transcriptomics findings with cardiac samples obtained from patients with HLHS and Tetralogy of Fallot from the database repository^29^. The single-cell analysis demonstrated that tissue samples from HLHS had an overall smaller number of cells including cardiomyocytes (CM), smooth muscle cells (SMC), and cardiac fibroblasts (CF) (Figure 6E) as compared to WT donor (Donor) and patients with tetralogy of Fallot (TOF). Comparing ratios of cardiac fibroblasts (CFs) with CMs and SMC demonstrated a higher portion of CFs as compared to CMs and SMCs in tissue from HLHS patients as compared to donor and patients TOF (Figure 6F), similar to ratios observed in our hiPSC CMs. In summary, CRISPR-edited NOTCH1 hypomorphs phenocopy CMs derived from the HLHS proband with NOTCH1 hypomorphism.

**Figure 6:**
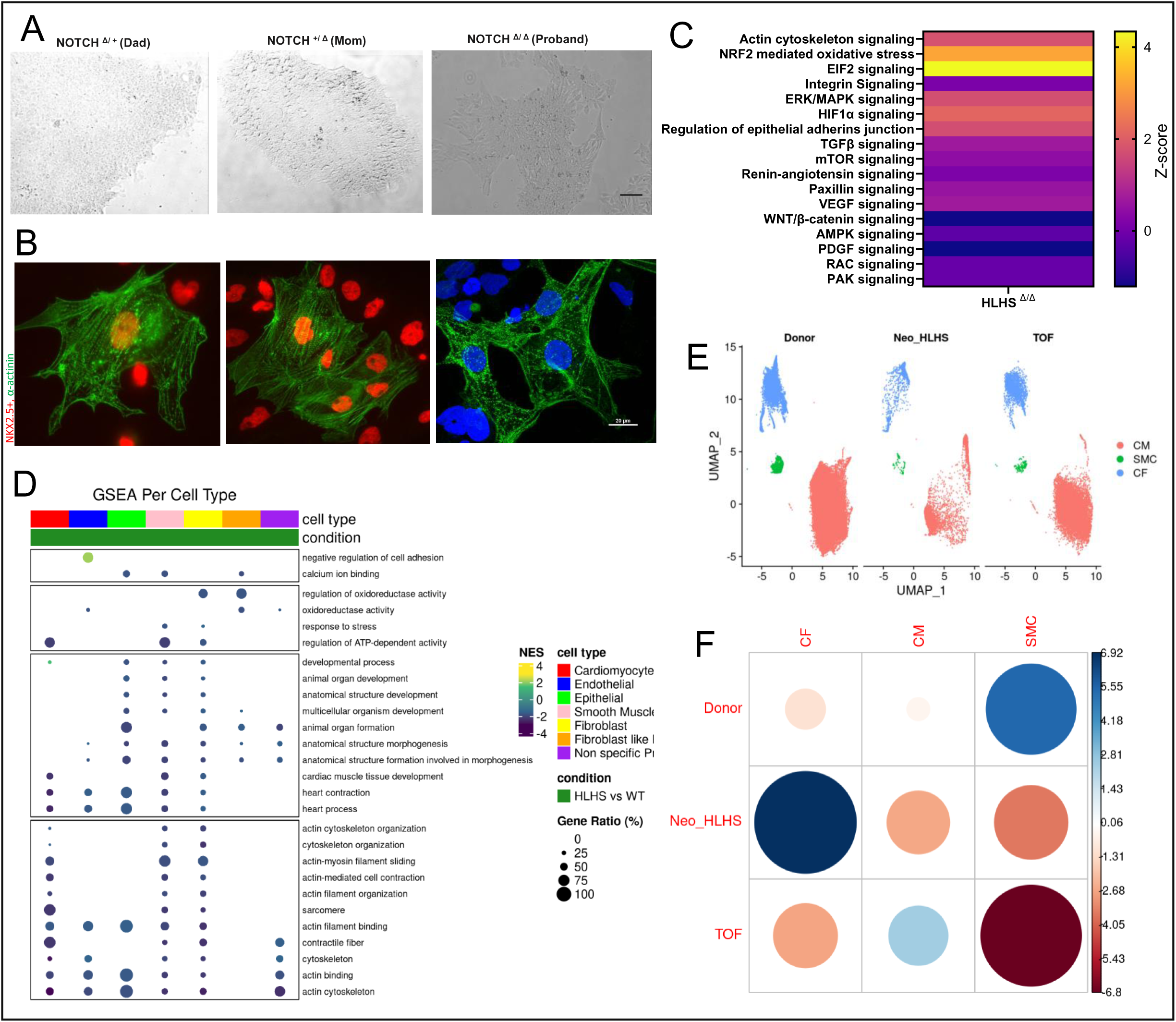
Patient-derived cells show a similar distribution as CRISPR-edited NOTCH1 hypomorphic iPS-CMs. (A) hiPSCs from proband with HLHS and non-HLHS parents show typical colony formation. Scale bar = 100µm (B) Cell differentiated into CM showed sparse and disorganized sarcomeric architecture for proband, whereas CMs obtained from the parents of the proband showed more dense and organized CM architecture (C) Differentially affected GO pathways demonstrated a similar dysregulation of the actin cytoskeleton, oxidative stress, and eIF2 signaling pathway for protein synthesis as seen in CRISPR-edited NOTCH1 hypomorphs (n=3 clones for each hiPSC donor). (D) GSEA per cell type analysis showed multiple pathways associated with cell development, cytoskeletal architecture, and electromechanical coupling were dysregulated, similar to those in CRISPR edited NOTCH1 hypomorphs (E) UMAP data from patient myocardial cells obtained from cardiac tissue of HLHS patients versus WT donor versus from patients with Tetralogy of Fallot showed an overall decrease in number of cardiac cells from HLHS patients (F) The relative abundance of CF cells were significantly higher in cardiac tissue from HLHS patients as compared to WT and TOF patients.

### High throughput proximity assay identifies auranofin as a positive regulator of NOTCH1 signaling

The preceding results demonstrate that the CRISPR-edited iPSCs faithfully recapitulate the disease state with respect to the morphological and signaling abnormalities seen in proband-derived cells and can be used as a potential platform for identifying therapeutics. To measure Notch1 ICD (NICD) protein level, a quantitative luminescence platform using the split Nano luciferase system was generated to allow high throughput assessment of NICD protein levels in cells. In brief, the NICD cDNA was cloned in-frame with a HiBiT tag, allowing for reconstitution with the rest of the Nanoluciferase enzyme (LgBiT) to provide a quantitative readout of NICD protein level (Figure 7A). HiBiT tagged NICD plasmid was transfected into BEAS-2B cells and stably transfected cells were plated onto a 384-well plate. A targeted subset of 250 FDA-approved drugs was screened in triplicate for increased luminescence after 18 hours of drug exposure (Figure 7B). The top hit, carfilzomib, a proteasomal inhibitor, was not selected for subsequent testing due to its known cardiotoxicity profile^30^ ^31^. The next most potent compound was auranofin, which has a known mechanism of inhibition of redox enzymes and is used for rheumatoid arthritis^32^. Significantly increased NICD protein level was noted in NICD-HiBiT expressing BEAS-2B cells following auranofin treatment (0.5µM, 18hr) (Figure 7C and 7D). Protein degradation using cycloheximide (CHX) chase assay demonstrated that auranofin stabilizes NICD, thereby delaying its degradation (Figure 7E). Increased endogenous NICD protein expression was also noted with auranofin exposure (100nM, 18 hours) in ECD^Δ/-^ hiPSCs (Figure 7F and 7G). A single-plate assay system was developed to screen the functional effects of auranofin on ECD^Δ/-^ hiPSCs (Figure 7I). Concentrations of auranofin between 5-400nM did not affect the viability of ECD^Δ/-^ hiPSCs cells. Increased and earlier beating was noted at concentrations between 5-100nM of auranofin (Figure 7J). In contrast, auranofin did not impact the beating, viability or NKX2.5 expression of WT hiPSC-CMs (supplemental figure 4). Immunofluorescence chemistry (IFC) of auranofin treated ECD^Δ/-^ iPSCs showed increased expression of NKX2.5+ cells (Figure 7K), with a significant increase in expression of NKX2.5+ as compared to untreated cells at 100 nM-300nM (31.86 ± 2.83 (0nM) vs. 61.00 ± 9.93 (100nM) vs. (61.67 ± 0.58 (200nM) vs. 54.67 ± 7.67 (300nM)) (Figure 7L). However, given that there was no beating observed in 200-300nM concentrations employed, 100nM of auranofin was used for subsequent testing. Brightfield analysis of beating ECD^Δ/-^ -CMs treated with auranofin versus vehicle controls showed a trend towards normalization of the beating waveform (Figure 7M), increase in amplitude of contraction, increase in beating frequency (decrease in peak time duration) and a decrease in duration of diastole and systole (Figure 7N). Auroanofin did increase NKX2.5 expression in HLHS proband cells suggesting that increase in NOTCH1 signaling was therapeutic in other cells with hypomorphic NOTCH1 expression (supplemental figure 4). Overall, auranofin was identified as a therapeutic target, that stabilized intracellular NOTCH1 expression and delayed degradation of NICD protein. Through these mechanisms, auranofin was able to improve NKX2.5 expression, and improve beating characteristics of hypomorphic NOTCH1 CMs in CRISPR edited and proband cells.

**Figure 7:**
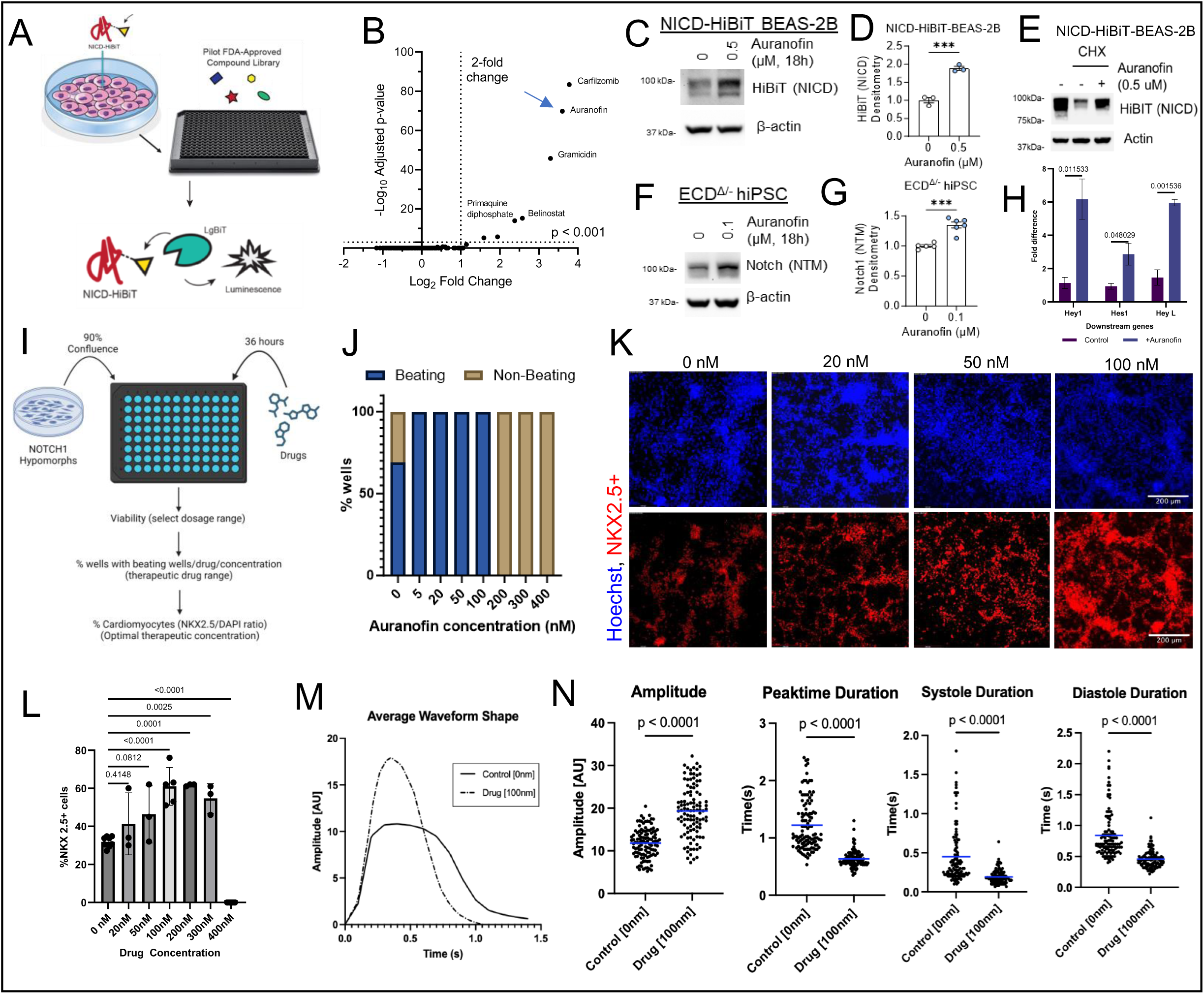
HiBiT technology-based screening assay identified Auranofin as a potential drug to facilitate CM differentiation in NOTCH1-HM-iPScs. (A) NICD-tagged HiBiT was stably expressed in BEAS-2B cells. A targeted subset of 250 FDA-approved drugs was screened for increased luciferase activity using a HiBiT assay. (B) Drugs that resulted in at least a 2-fold increase in luciferase activity were screened as positive candidates for further screening. Auranofin, identified through the screen, was tested for improved cardiomyocyte differentiation in subsequent assays (C) Western blot showed increased expression of HiBiT tagged NICD with 500nM of Auranofin on BEAS-2B cells (D) with a 2-fold increase in expression. (n= 3). (E) Treatment of BEAS-2B cells without and with cycloheximide showed increased degradation of HiBiT-NICD respectively; however, this degradation rate was decreased in the presence of Auranofin, suggesting that Auranofin acts through post-translational stabilization of the protein. (F) Treatment of ECD^Δ/-^ hiPSCs with Auranofin similarly showed an increased expression of NICD by a (G)1.4-fold (n=5-6). (H) Addition of Auranofin to ECD^Δ/-^ hiPSCs showed an increase in target gene expression of Hey1, Hes1 and HeyL. (n=3 clones) (I) A single plate assay was developed to test drug candidates’ efficacy. NOTCH1 hypomorphs were plated on a 96-well plate and treated with an increasing dosage of Auranofin for up to 36h. (J) hiPSCs were differentiated (n=3) and analyzed 15 days post- differentiation. Wells were screened for cell loss (indicative of toxic drug concentrations), and the presence of beating CMs, which was found to be increased at Auranofin concentrations of 5-100 nM. (K) Wells were then stained for CM marker NKX2.5 using IFC. (L) Compared to the vehicle alone, Auranofin 5nM-100nM showed an increase in NKX2.5+ expression between 200-400nM. NKX2.5 positive cells were significantly increased with 100nM of Auranofin (n=3, p<0.0001). In aggregate 100nM of Auranofin showed both an increase in beating CMs and NKX2.5 expression (n=3). (M) The average waveform of beating cells analyzed in brightfield imaging showed improved waveform characteristics with a similar waveform to WT cells. (N) Wave amplitude increased, peak time and duration of systole and diastole decreased, indicating improved beating characteristics. (n=107-114 cells from 1-2 organoids per 3 clonal replicates).

## Discussion

This study describes the creation of a collection of isogenic iPSCs with hypomorphic NOTCH1 expression by CRISPR-Cas9-medited editing. These iPSCs have distinct transcriptomics profiles both as iPSCs and differentiated cardiac organoids, as well as distinct cellular and functional properties. Given the clinical relevance of these mutations, and the ability of this cellular model to recapitulate various aspects of CHD, we were able to use these cells to validate novel agents that might have therapeutic value in HLHS patients.

The genetics of CHD are complex and likely contribute to the variable penetrance seen in CHD family cohorts^33^. This complex genetic pattern of disease contributes to the difficulty in generating animal models that recapitulate the disease. In that context, hiPSC-based in vitro cellular systems can provide significant insight into genotype/ phenotype correlations of this complex disease. Our studies, using robust isogenic controls, support observations seen by others where iPSCs derived from an HLHS proband-parent trio demonstrated increased disorganization of myofibrils in the HLHS proband compared to the parental cells^20^. In this case, the proband represents a compound heterozygote inheriting a mutant NOTCH1 allele with an extracellular mutation in NOTCH1 from the father, and a NOTCH1 mutant allele with an intracellular mutation from the mother. While neither parent had HLHS, the mother did have a bicuspid pulmonary valve, while the father had no structural heart disease. The parent’s phenotype suggests that hypomorphic mutations affecting different regions of NOTCH1 may have different clinical presentations. To further pursue this notion, we generated hypomorphic mutations in either the extracellular or intracellular domains of NOTCH1, to determine the impact of disrupting different aspects of NOTCH1 signaling on CM physiology. While both the ECD and ICD domains contributed significantly to abnormalities in pluripotency and CM differentiation, significant disruption was noted in cytoskeletal signaling pathways associated with actin, both pre- and post-differentiation.

NOTCH1 is a known mechano-transduction protein that has been implicated in arterial endothelial cells in shear stress responses^34^. However, its role in cardiomyocytes is not as well understood. Our studies demonstrate that multiple proteins involved in cytoskeletal signaling and mechano-transduction are dysregulated in the pre-differentiated hiPSCs. hiPSCs show a significant dysregulation of actin cytoskeletal proteins including α-actinin (ACTN2 and ACTC1) which are essential in the formation of z-discs in CM sarcomeres. Proteins interacting with the cytoskeleton, as well as integrins, are also dysregulated including paxillin (Pak1), microtubule- associated protein 2 (MAP2), TRIO and F-actin binding protein (TRIOBP), and Ankyrin 2 (ANK2). Additionally, proteins that facilitate cell adhesion and communication (GJA1, CAV1) are also disrupted. Significant abnormalities in cytoskeletal signaling proteins in the pre-differentiation stage likely contribute to subsequent abnormalities in structural proteins when cells differentiate into cardiomyocytes. The abnormalities are demonstrated through our transcriptomics studies and have previously been demonstrated by other groups in their HLHS CMs studies^25^.

Abnormalities in iPSCs cytoskeletal signaling lead to an abnormal cytoskeleton as noted in ours and other studies, with a significant overlap of phenotype between CM, FB, and SCM. Functionally, this translates to a decrease in calcium transients and contractility. This change in morphology is also observed in the HLHS proband as well as myocardial tissue from HLHS patients. As mitochondria provide ATP for sarcomere contraction and develop synchronously as previously demonstrated^35^, mitochondrial density directly correlates to Z-disc cross-sectional area. NOTCH1 hypomorphic clone-CMs show a similarly disrupted mitochondrial pattern of distribution, decreased ATP production, and increased ROS production.

The close correlation of NOTCH1 hypomorphic clones generated through CRISPR gene editing with both HLHS patient-derived iPSC-CMs and myocardial tissue obtained from HLHS patients confirmed the clinical validity of using these HC clones for assessing therapeutic candidates. We demonstrate here that some currently available FDA-approved agents can regulate NOTCH1 ICD levels. We used a heterologous HiBiT reporter cell line to first identify potential hit molecules due to the difficulty of using iPSCs in high-throughput screens. Potential hits were then individually assessed with our hypomorphic clones for their ability to modulate endogenous NOTCH1 ICD, and to rescue the various defects in differentiation and function seen with our hypomorphic NOTCH1 clones. The ability of Auranofin to revert NOTCH1 hypomorphic clones to a more wild-type phenotype identifies this agent as a potential therapeutic intervention. The sub- micromolar concentrations required in vitro to restore CM function are well within the therapeutic plasma levels of this agent when it is employed for rheumatoid arthritis^32^. Of note, auranofin is currently being explored for numerous other disease states besides arthritis including infectious diseases and cancer^36^. While its precise mechanism of action is unclear, some evidence suggests that it may act as a proteasomal inhibitor, albeit working differently than agents such as bortezomib^30^. This mechanism of action is consistent with the top hit molecule from the screen which was the known proteasomal inhibitor Carfilzomib. Given that the NOTCH1 ICD is actively regulated by the ubiquitin-proteasomal system^37,38,39^ agents that perturb this system might have beneficial effects. While we concentrated our efforts on FDA-approved agents due to their ability to be rapidly re-purposed, the platform described could be easily adapted to help identify novel therapeutic agents that might have even more desirable effects.

### Limitations of the study

NOTCH1 is implicated in approximately 8% of known patients with HLHS and other left-sided congenital cardiac lesions. Hence, other genetic mutations and interactions may be involved in patients with HLHS and hence have other signaling pathways implicated in their pathology. We have primarily used hiPSCs and human tissue data to demonstrate and confirm our findings. There are no known animal models that involve complex interactions between various genes and hence could not be used here. Lastly, we identified and characterized Auranofin as a potential therapeutic topic. While Auranofin has numerous beneficial targets, its off-target liabilities and translational ability will need to be further studied.

## Acknowledgments

AS and the experiments described here are supported by the American Heart Association Career Development Award (852875) and the NHLBI K08 161440. The drug discovery aspect is supported by the Heart Fest Grant for Congenital Heart disease. TL is funded by K99 AG078342. JL is supported by the Clinical Scientist Training Program. Transcriptomics analysis was supported in part using the HTC cluster, which is supported by NIH award number S10OD028483 and internal funds from the Children’s Hospital of Pittsburgh.

## Supplemental Figures and Legend

**Supplemental Figure 1:**
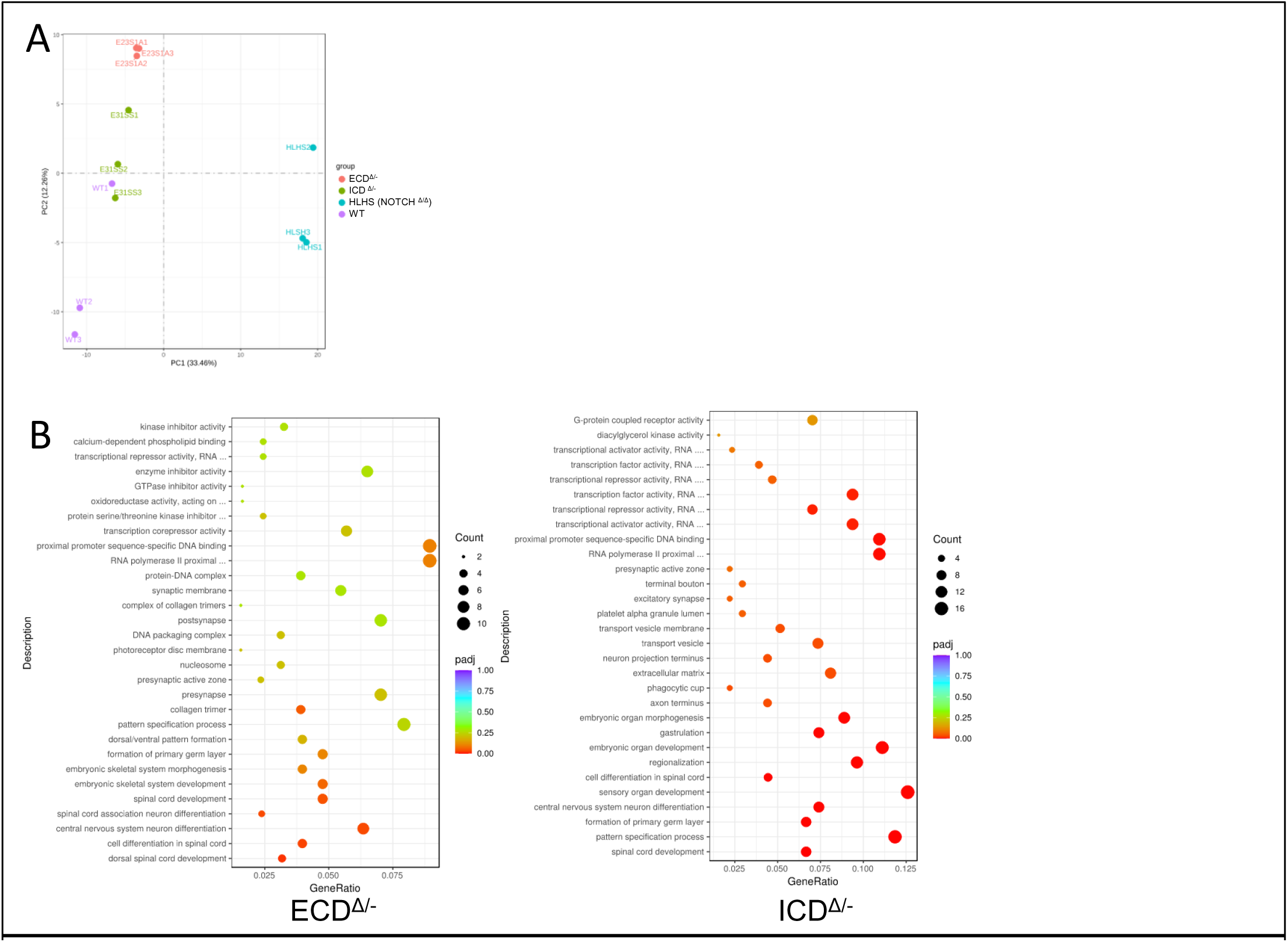
(A) Principal component analysis of transcriptome clusters of WT versus ECD ^Δ/-^ versus ICD^Δ/-^ versus HLHS proband as compared to WT hiPSC. (n=3 clones per condition). (B) GO enrichment analysis scatter plot of ECD^Δ/-^ vs. ICD^Δ/-^ shows similarities and differences in signaling pathways differentially regulated with ECD versus ICD domain mutations in the iPSC pre-differentiation stage. While both loci of mutations result in dysregulation of important pathways associated with protein synthesis, pathways associated with oxidative activity and calcium dependent signaling are affected in ECD^Δ/-^ versus ICD^Δ/-^ iPSC clones.

**Supplemental Figure 2:**
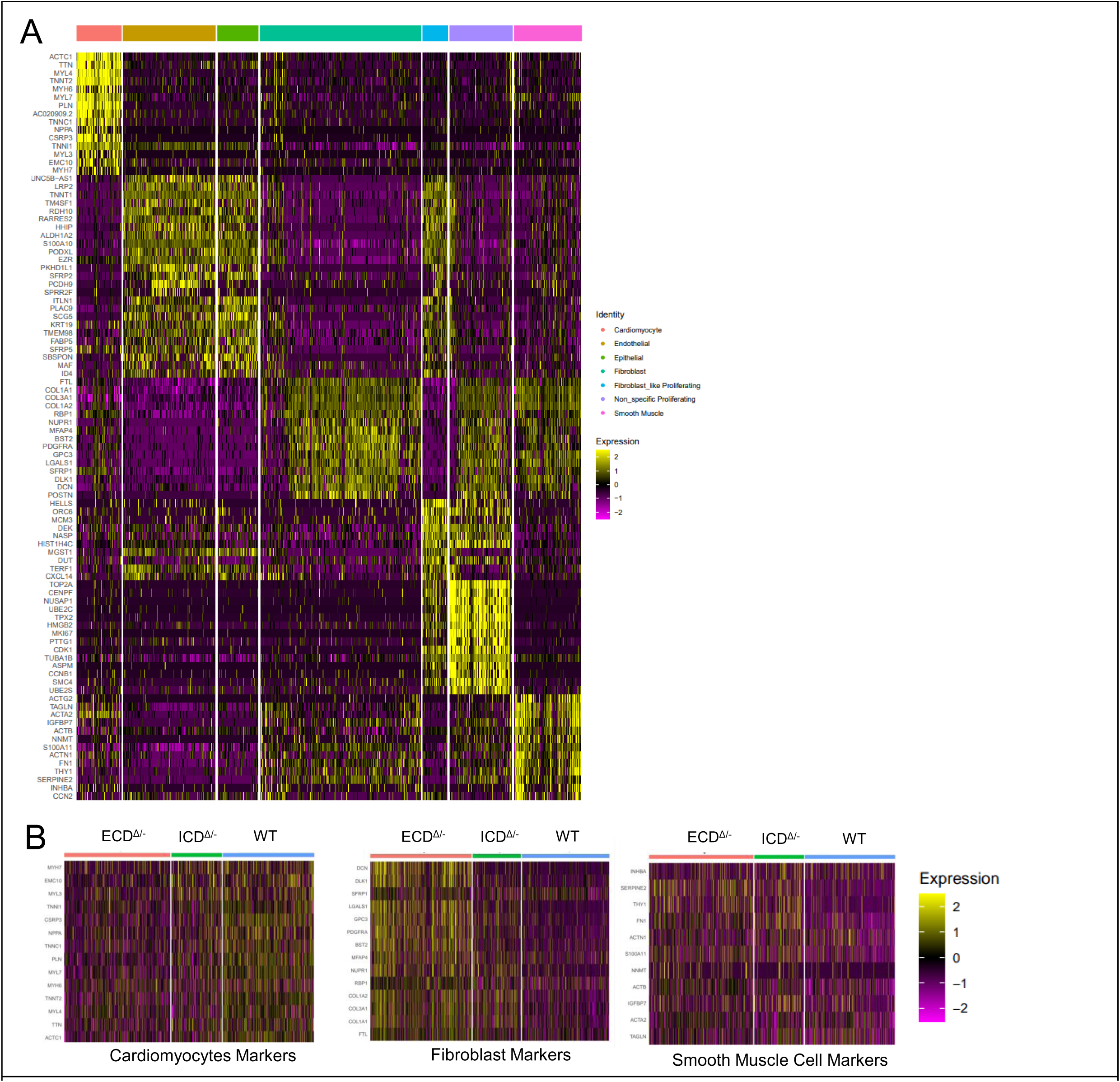
(A) Heat Map of cell-specific markers across the various cell types: Cardiomyocytes (CM), endothelial cells (EDC), epithelial cells, fibroblasts, fibroblast-like proliferating, non-specific proliferating and smooth muscle cells. (B) Heat-map of cell-specific markers associated with cardiomyocytes, fibroblasts and smooth muscle cells show that while

**Supplemental Figure 3:**
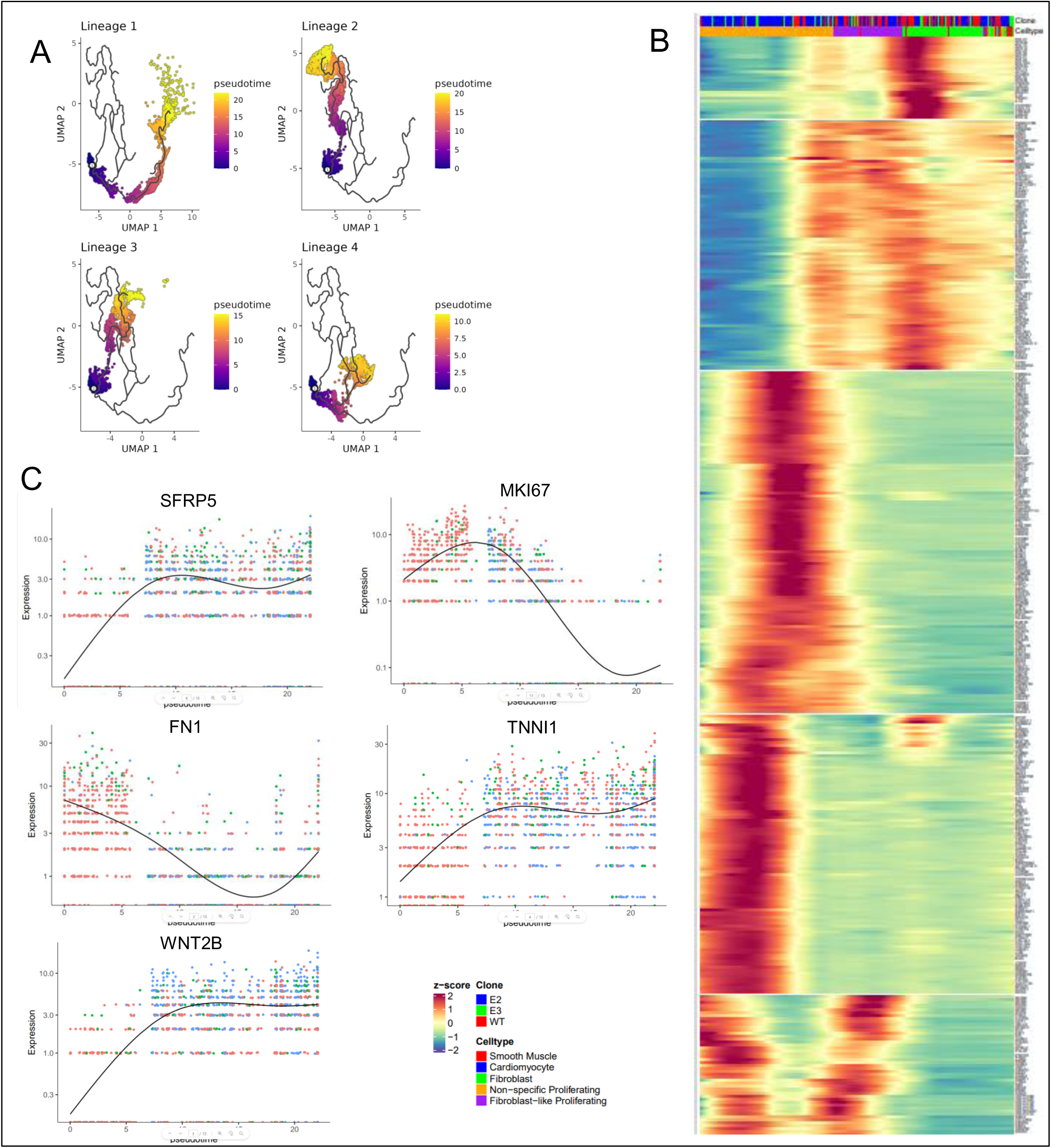
(A) Four lineages were identified as followed by various cell subtypes (B) Pseudo time analysis of lineage 1 shows a . z-score transformed expression profiles along Lineage1 trajectory where Y-axis represents the various genes surveyed and X-axis represents the pseudo time in which the cells were ordered. (C) Expression profiles of representative genes of the major modules

**Supplemental Figure 4:**
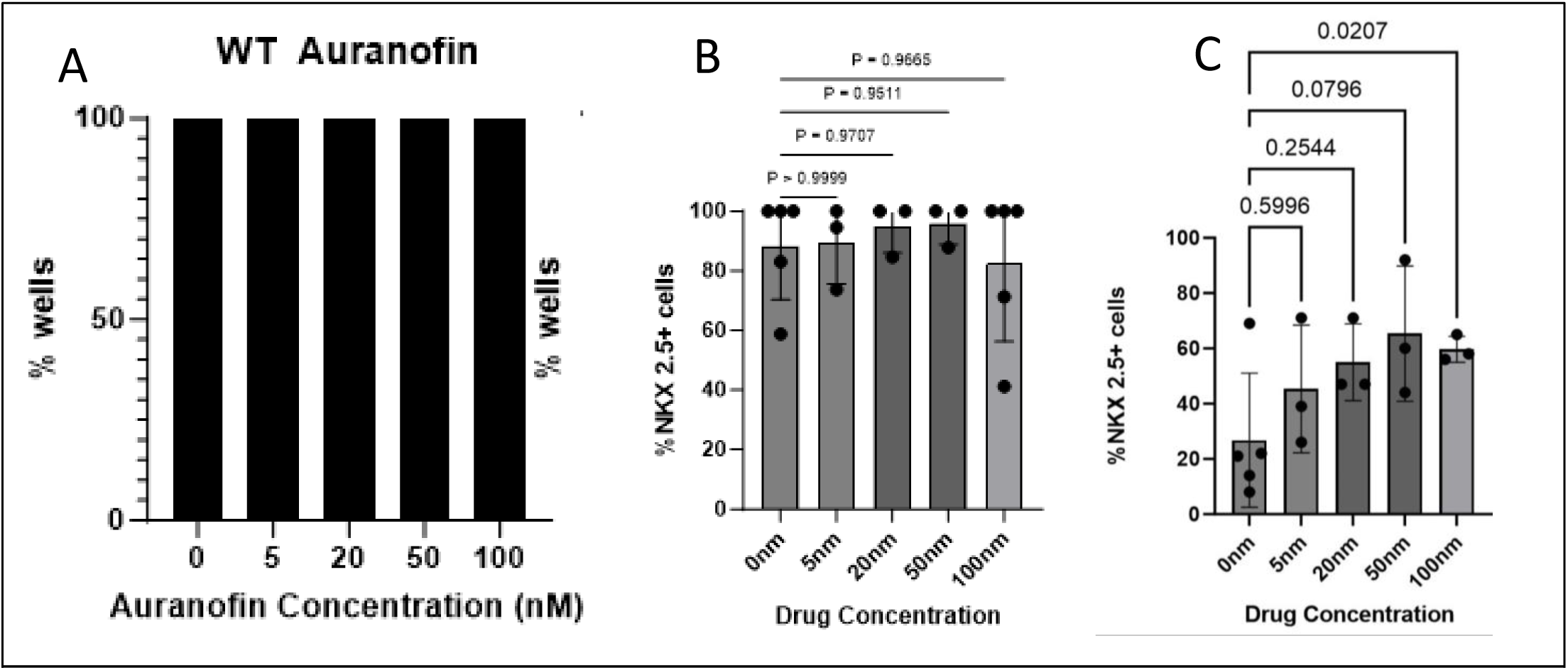
(A) WT iPSCs treated with auranofin (concentration ranges between 0-100nM) all demonstrated normal beating patterns indicating no toxic effects of auranofin on WT cells. (B) NKX2.5 staining showed no significant decrease CM differentiation with auranofin (n=3 wells per condition with multiple regions of interest per well). (C) HLHS Patient-derived iPSCs with hypomorphic NOTCH1 expression showed a significant increase in NKX2.5 expression at 100 nM auranofin concentration. (n=3 independent)

## Notes

### Competing Interest Statement

The authors have declared no competing interest.

