## supplemental for "Hypomorphic NOTCH1 Expression Alters Cardiomyocyte Cellular Architecture in Hypoplastic Left Heart Syndrome"

### Key Resource Table

| REAGENT or RESOURCE | SOURCE | IDENTIFIER |
|---|---|---|
| **Antibodies** | | |
| Antibody against HiBiT | Promega | Cat# 30E5, RRID:AB_2924793 |
| Antibody against Notch1 (WB) | Cell Signaling Technology | Cat# 3608, RRID:AB_2153354) |
| Antibody against Notch1 (IFC) | Invitrogen | Cat # MA5-11961 |
| Antibody against NKX2.5 | Life Technologies (Fisher) | Cat# 710634 |
| Anti-α-Actinin | Sigma | A7811 |
| Antibody-goat-anti-mouse-Alexa Fluor 488 | Invitrogen | A21121 |
| Antibody-goat-anti-rabbit- Alexa Fluor 594 | Invitrogen | A-11012 |
| Antibody-donkey-anti-mouse-Alexa fluor 594 | Invitrogen | A21203 |
| Enzymes and buffers | | |
| T7 Endonuclease I | NEB | Cat# M0302L |
| BsmBI | NEB | Cat# R0580 |
| NEbuffer 3.1 | NEB | n/a |
| NEbuffer 2 | NEB | n/a |
| BbsI-HF | NEB | Cat# R3539S |
| CutSmart | NEB | n/a |
| EcoRI | NEB | Cat# R3101S |
| **Plasmids** | | |
| pFC37K-HiBiT plasmid | Promega | Cat# N2391 |
| pCHD-CMV | Addgene | Plasmid #72265 |
| ACTN2-pEGFP | Addgene | Plasmid # 52669 |
| lenti-CRISPR_v2 | Addgene | Plasmid # 98290 |
| pU6-pegRNA-GG-acceptor | Addgene | Plasmid #132777 |
| pCMV-PE2 | Addgene | Plasmid # 132775 |
| **Guide RNA** | | |
| gRNA1 for ECD$^{\Delta/-}$ | Sigma | 5’ CCGGGCTTCGTGGGTGAGCG 3’<br>5’ CGCTCACCCACGAAGCCCGG 3’ |
| gRNA2 for ECD$^{\Delta/-}$ | Sigma | 5’GGCTTCGTGGGTGAGCGCTG3’<br>5’CAGCGCTCACCCACGAAGCC3’ |
| gRNA1 for ICD$^{\Delta/-}$ | Sigma | 5’GGGCAACAGCGAGGAAGAGG3’<br>5’CCAAGCGCCTGCTGGAGGCC3’ |
| gRNA 2 for ECD$^{\Delta/-}$ | Sigma | 5’CCAAGCGCCTGCTGGAGGCC3’<br>5’GGCCTCCAGCAGGCGCTTGG3’ |
| **Chemicals, peptides and recombinant proteins** | | |
| B-27™ Supplement, minus insulin | ThermoFisher Scientific | Cat# A1895601 |

| | | |
|---|---|---|
| B-27™ Supplement(50X), serum-free | ThermoFisher Scientific | Cat# 17504044 |
| Roswell Park Memorial Institute (RPMI) 1640 medium | ThermoFisher Scientific | Cat# 11875093 |
| DMEM F12 | ThermoFisher Scientific | Cat# 11320033 |
| FBS | Life Technologies (Fisher) | Cat# 16000036 |
| Hoechst | Life Technologies (Fisher) | Cat# H1399 |
| Matrigel | Corning (Fisher) | Cat# CB40234 |
| PBS | ThermoFisher Scientific | Cat# 20012050 |
| EDTA 50nM | ThermoFisher Scientific | Cat# AM9260G |
| E8 Basal Media | Life Technologies | Cat# A1517001 |
| Opti-MEM I Reduced Serum Medium | Thermo Fisher Scientific | Cat # 31985070 |
| Lipofectamine Stem Transfection Reagent | Thermo Fisher Scientific | Cat # STEM00001 |
| Trypsin | ThermoFisher Scientific | Cat# 25-200-056 |
| Fluo-4 AM Cell permeant 10x50ug | Life Technologies (Fisher) | Cat #F14201 |
| Mitotracker Red CMX ROS 20x50ug | Life Technologies (Fisher) | Cat# M7512 |
| Y-27632 Dihydrochloride (ROCK inhibitor) | Santa Cruz | Cat# SC-281642A |
| iPSC Cardiomyocyte Differentiation Kit | Life Technologies (Fisher) | Cat# A2921201 |
| Oligomycin | Sigma | 75351 |
| FCCP | Tocris | 0453 |
| Rotenone | Cayman | 13995 |
| Antimycin A | Sigma | A8674 |
| Rhod-2, AM | ThermoFisher Scientific | Cat# R1244 |
| **Cell Sources** | | |
| hiPSCs | National Institutes of Health commonfund.nih.gov/stemcells/lines | ND2.0 |
| HLHS trio hiPSC | Mayo Clinic, Rochester, MN | Materials Transfer Agreement |
| HEK293T | ATCC | Cat # CRL-3216 |
| BEAS-2B | ATCC | Cat # CRL-3588 |
| **Primers for rt-PCR** | | |
| Hey2-F | Sigma | TTGAAGATGCTTCAGGCAACAGGG |
| Hey2-R | Sigma | TCAGGTACCGCGCAACTTCTGTTA |

| | | |
|---|---|---|
| Hey1-F | Sigma | AGAGTGCGGACGAGGAATGGAAACT |
| Hey1-R | Sigma | CGTCGGCGCTTCTCAATTATTCCT |
| Notch1-F | Sigma | CGCACAAGGTGTCTTCCAG |
| Notch1-R | Sigma | AGGATCAGTGGCGTCGTG |
| Hes1-F | Sigma | AGTGAAGCACCTCCGGAAC |
| Hes1-R | Sigma | TCACCTCGTTCATGCACTC |
| KLF4 F | Sigma | ACGATCGTGGCCCCGGAAAAGGACC |
| KLF4 R | Sigma | TGATTGTAGTGCTTTCTGGCTGGGCTCC |
| OCT3/4F | Sigma | GACAGGGGGAGGGGAGGAGCTAGG |
| OCT3/4 R | Sigma | CTTCCCTCCAACCAGTTGCCCCAAAC |
| SOX2 F | Sigma | GGGAAATGGGAGGGGTGCAAAAGAGG |
| SOX2 R | Sigma | TTGCGTGAGTGTGGATGGGATTGGTG |
| MYC F | Sigma | GCGTCTGGGAAGGGAGATCCGGGC |
| MYC R | Sigma | TTGAGGGGCATCGTCGCGGGAGGCTG |
| GAPDH F | Sigma | GGACCTGACCTGCCGTCTAGAA |
| GAPDH R | Sigma | GGTGTCGCTGTTGAAGTCAGAG |
| **Kits** | | |
| CellTiter-Glo | Promega | Cat # G9241 |
| Nano-Glo HiBiT Lytic Reagent | Promega | Cat # N3030 |
| Plasmid mini-prep kit | Zymogen | Cat # D4209 |
| Quick RNA-miniprep Kit | Zymogen | Cat # Z5223 |
| Gel DNA recovery Kit | Zymogen | Cat # D4001 |
| Phusion HF polymerase Kit | Thermo Fisher | Cat # F531S |
| Seahorse XF96 Cell Culture Microplate | Agilent Technologies | Cat # 101085-004 |
| **Software and algorithms** | | |
| Image J | NIH | https://imagej.net/Fiji/Downloads |
| CRISPR sgDesign | Broad Institute | portals.broadinstitute.org/gpp/public/analysis-tools/sgrna-design |
| GraphPad Prism10 | GraphPad | N/A |
| Leica Application Suite X software | Leica | N/A |
| Spiky | GitHub | https://github.com/PCCV/Spiky |
| Myocyter | GitHub | https://github.com/Myocyter/Myocyter-v1.3 |
| Drug Library | TargetMol | https://www.targetmol.com |

**Method details:**

**DNA cloning:**

**Synthesis of CRISPR-Cas9 vectors:** CRISPR-gene editing was used to create mutations in the ECD and ICD loci in exon 23 and exon 31 respectively using two sgRNAs per loci. The sgRNA were designed using the sgRNA design tool from The Broad Institute, MA. lentiCRISPRv2 puro was a gift from Brett Stringer (Addgene plasmid # 98290)[40]. Additionally, pU6-pegRNA-GG-acceptor and pCMV-PE2 were a gift from David Liu (Addgene plasmid # 132777) and (Plasmid # 132775)[41]. Plasmids generated from sgRNA design tool were synthesized and cloned into lentiCRISPRv2 with a sequential double digest. sgRNA integration was confirmed using PCR and Phusion HF DNA polymerase kit.

**Synthesis of pCHD-ACTN2-eGFP plasmid constructs:** ACTN2 was cloned into the pCDH-CMV-MCS, between restriction sites BamHI and NotI, with a GSGSGS peptide linker sequence. For construct assembly, the pCDH-CMV-MCS vector (50 ng), and the corresponding ACTN2-eGFP fragment with linker sequences was incubated at 50˚C for 15 mins after which it was transformed into bacteria and amplified overnight in LB. Plasmids were purified using DNA miniprep.

**Synthesis of HiBit Tagged NICD constructs:** HiBiT-tagged NICD plasmid constructs were generated using molecular cloning and the FLEXI system (Promega). Briefly, the open reading frame of the target gene was PCR amplified with restriction sites for AsiSI and PmeI and was cloned into pFC37K-HiBiT plasmid. NICD template was generated from TetO-FUW-NICD, which was a gift from Rudolf Jaenisch (Addgene plasmid #61540)[42].

All plasmid constructs were verified by DNA sequencing (Genewiz).

**Culture and expansion of iPSCs:**

Wild-type hiPSCs were obtained from the NIH iPSC repository (ND2.0, NIH CRM control iPSC line) and plated on Matrigel-coated plates and supplemented with E8 supplemental medium until they reached 80-90% confluence in a 12-well plate. Cells were split at least 3 times using EDTA, at 80-90% confluence at which time cells were at which point they were split and transferred to a 12-well plate. iPSC colonies were closely monitored for aggregation and phenotype and cells with spindle-shaped morphology and spindle-shaped cells were excluded in subsequent cultures by selecting rounded colonies.

**Generation of isogenic hiPSC hypomorphic NOTCH1 clones:**

Protocol is modified from WT iPSCs are plated onto a 12-well plate overnight to achieve <25% confluence. Before transfection, Opti-MEM I medium and Lipofectamine Stem Transfection Reagent were warmed to room temperature. hiPSCs with E8 medium containing were pretreated with ROCK inhibitor for 30 min in the 37°C incubator. Approximately 2 µg of plasmid per reaction was aliquoted into a microcentrifuge tube. 50µl Opti-MEM I per reaction was added to each tube together with 12.5 µl Lipofectamine Stem Transfection Reagent per reaction. The reaction was gently mixed and incubated for

10 min at room temperature. The complexes were then distributed to the wells of pretreated hiPSCs cells. Cells were incubated for 48 hours after which they were treated with 0.5 μg/mL of Puromycin for a period of 5-7 days. Media was changed every 48 hours and fresh E8 medium with Puromycin was added. Individual colonies of cells were picked under a microscope using a sterile pipette tip and transferred to a fresh 12-well plate where independent clones were transferred to their own well. Independent clones of cells were allowed to expand in the presence of Puromycin, after which they were split and expanded again. A badge of CRISPR-edited clones were then lysed for sequencing (GeneWiz) and their corresponding clonal expansions were frozen for further use if sequencing confirmed the intended DNA modifications. Off-target effects were checked.

**Transfecting live CMs with GFP-tagged α-actinin**

WT and NOTCH1 hypomorphic hiPSCs were differentiated into CMs. At day 15 post differentiation, pCDH-CMV-ACTN2-EGFP plasmids were transduced into beating CMs in the presence of Lipofectamine 200 and after 48 hours of exposure the supernatant was exchanged with cell culture medium. Beating cells were imaged on a Nikon Tie 8200, NY in a custom-made humidity, oxygen, and temperature-regulated stage at 37˚C

**iPSC differentiation:**

WT and CRISPR-edited hiPSCs were differentiated in 12 well plates, after reaching 90% confluence using Cardiomyocyte differentiation media A and B per protocol. When organoids were generated, cells were transferred to an AggreWell plate at day 7 post differentiation, after which the plate was centrifuged at 100 rpm at 10 mins and incubated at 37˚Celsius for 48 hours. Subsequently, organoids were dislodged from the AggreWell plate and transferred into non-adherent plates maintained in RPMI B27+ insulin media.

**Mitochondrial Analysis**

Immunofluorescence was performed to visualize mitochondrial networks using Mitotracker red. Each sample was imaged at 100x in sections and stitched together using Image J. To analyze differences among mitochondria within the different cell lines, we utilized a Fiji plugin; Mitochondrial Analyzer. This tool quantifies and qualifies mitochondrial network morphology and dynamics by binarizing images based on adaptive thresholding methods. This identifies mitochondrial objects which are then analyzed in 2D to provide quantities for morphological parameters including mitochondrial count, area, perimeter, form factor, and aspect ratio. The identified mitochondrial objects are then skeletonized and analyzed to quantify the number of networks and degree of branching (number of branches, length of branches, number of branch junctions). Once uploaded into Image J, images were converted to 8-bit. Pre-analysis processing was done through the Mitochondrial Analyzer interface. The method used to determine the adaptive threshold was the weighted mean.

Optimization of the 2D threshold was achieved with a block size of 1.65 and a C-value of 5. Default settings of the program were used, and analysis was performed on a per-cell basis and network characteristics were normalized to the mitochondrial count. Statistical analysis was performed using one-way ANOVA followed by Tukey's post-hoc test.

**Seahorse assay:**
20,000 iPSC-CMs were seeded in a 96-well plate and assayed for oxygen consumption rates (OCR) using a Seahorse XFe96 flux analyzer (Agilent). Following measurements of basal OCR, samples were injected with 1 μM of oligomycin, an ATP synthase inhibitor to measure ATP-linked respiration and proton leak. This was followed by 0.5 μM carbonyl cyanide-4-(trifluoromethoxy)-phenylhydrazone (FCCP) injection to determine maximal OCR and the spare respiratory capacity of the cells. All cells were finally injected with Antimycin-A-Rotenone (1 μM each) to completely inhibit the electron transport chain and measure non-mitochondrial oxygen consumption rates.

**Contractility Analysis**
Brightfield videos of unpaced, spontaneously beating cardiomyocytes at 37˚C were recorded at 20x for 15 to 30 seconds. Beat frequencies were manually calculated and outliers were excluded from analysis. Twenty-four samples (n=10 WT, 7 $ECD^{\Delta/-}$, 7 $ICD^{\Delta/-}$) were included in the analysis with additional exclusion criteria for analysis based on cell visibility and video quality, such as blurriness, artifacts, shakiness, and inconsistent brightness. To analyze differences among contractility within the different cell lines, we utilized an Image J macro, Myocyter. This tool characterizes contracting regions using two parameters: "speed," which is the difference between consecutive frames of a video and indicates the speed of a contraction, and "amplitude," which is the difference between the current frame and an automatically determined reference frame of the cell in its resting phase and indicates the amount of deformation of the contracting cell compared to the reference frame. The difference between the images is then calculated and presented as a plot and numerical output, generating values for mean frequency, amplitude, systole (contraction time), diastole (relaxation time), peak times (total length of contraction and relaxation), and beat time (peak to peak time). Using dynamic thresholding, the program can precisely track amplitudes even when the baseline shifts. Once uploaded into Image J, the "ROI (Region of Interest) Manager" tool was utilized to select contracting cells within each video, with approximately 15 to 40 cells identified per video. Next, the Myocyter macro was run using the manual ROI list and default settings for intensity threshold, particle size, peak recognition, and sensitivity of maxima and minima detection. Analysis was performed on a per-cell basis, and graphical results were evaluated for quality control (correct detection of beats). Statistical analysis was performed using one-way ANOVA followed by Tukey's

post-hoc test.

**Calcium Fluorescence Analysis**

Videos of unpaced, spontaneously beating cardiomyocytes at 37˚C were captured at 20x using fluorescence imaging. To visualize calcium transients within cells, calcium indicator Fluo-4 was used at excitation 488nm and emission wavelength 509nm. Images with debris and videos with <50 fps were excluded from the analysis, which resulted in n=10 WT, 7 ECD$^{\Delta/-}$, and 8 ICD$^{\Delta/-}$ for final analysis. Calcium transients were quantified using an Image J plugin, Spiky, which quantified the amplitude, time to peak, time between peaks, full and half-width at 50% maximum, slope of peak, and area under the curve of the calcium transient.  Approximately 40-50 ROIs were identified per video. Graphical results were evaluated for the correct detection of peaks and those samples with errors were either corrected or excluded from analysis. Statistical analysis was performed using one-way ANOVA followed by Tukey's post-hoc test for the parameters evaluated.

**Analysis of sarcomeres:**

The sarcomeric organization was characterized by identifying 50 CMs per group, by three independent readers blinded to the control and experimental groups. 1 point was assigned to the presence of filamentous cytoskeletal structures, 1 point was assigned for the presence of organized sarcomeres, and one point was assigned to the presence of z-discs throughout the sarcomeres for a maximum total of 3 points. The scores were averaged across the three readers. Hence, overall scores were classified where Class I contained unassembled or newly formed sarcomeres, being sparse, irregular, with gaps, and not yet directionally organized; Class II corresponded to sarcomeres showing higher density and more regular space, but which only just started to be directionally organized; class three contained linear and directionally organized sarcomeres in which Z-disk banding is not yet distinct or predominant, and Class III corresponded to developed sarcomeres with established directionality and running in parallel, with clear banding of Z-disks

**Processing and analysis of hiPSC transcriptomics**

tRNA from undifferentiated hiPSCs of WT and NOTCH1 hypomorphs was extracted and sent to Novogene Inc for transcriptomics analysis. Following receipt, the tRNA was checked for quality control, following which the samples were analyzed, and the RNA library was constructed and sequenced. Data analysis was performed by Novogene, Inc. For the generation of cord diagrams, an R-package (Circlize) was used. Data provided by Novogene related to gene expression was used to create heat maps as well as Z-scores associated with the expression of the hypomorphic clones as compared to WT cells.

**Pre-processing and alignment of scRNA-seq data**

Three replicates each of CMs from WT, ECD$^{\Delta/-,}$ ICD$^{\Delta/-,}$ and HLHS samples were multiplexed according to the 10X CellPlex protocol and sequenced. Resultant bcl files were demultiplexed using the bcl2fastq utility from 10X, generating FASTQ files for each set of samples (WT, ECDΔ/-, ICDΔ/-, and HLHS). These files were then processed using the CellRanger (V6.1.1)[43] "multi" protocol for alignment against the GRCh38 human genome (refdata-gex-GRCh38-2020-A) using GEX and Multiplexing Capture assays according to the standard CellRanger "multi" workflow. This resulted in a set of per-sample gene-count matrices, demultiplexed for downstream analysis and processing.

**Quality control:**

We implemented a standard Seurat (V4.3.0)[44] based workflow to pre-process the sample datasets. Pre-processing steps included merging the replicates, with quality control being performed at both the cell and gene level. We excluded empty cells, doublets, and possible multiplets from the analysis, by determining a gene count threshold based on the median number of genes using quality control violin plots and heatmaps. Cells were filtered based on a minimum number of 200 genes per cell as well as a mitochondrial gene percentage threshold of 15% to exclude dead or dying cells. After QC, 10916 high-quality cells were selected for further analysis. The HLHS samples were later removed, which resulted in a set of 9209 high-quality cells.

**Single-cell integration and clustering**

The biological replicates for each condition (WT, ECD$^{\Delta/-,}$ ICD $^{\Delta/-}$) and the data from corresponding samples were analyzed using a standard Seurat integrative data analysis pipeline. The FindVariableFeatures function was used to select the top highly variable integration anchors, which were then subjected to a Principal Component Analysis (PCA). Robust bootstrapping methods such as the jackstraw procedure and elbow heuristics were employed to identify 15 significant principal components, which were then used as the input for the Uniform Manifold Approximation and Projection (UMAP) dimensionality reduction. Nearest-neighbors and clusters were derived from the UMAP graph using the standard Seurat shared nearest-neighbor calculations (FindNeigbors and FindClusters), with a clustering resolution of 0.5.

**Annotation of unique cell clusters**

For cell type annotation, we calculated the top expressed marker genes for each cluster using a one-versus-all differential expression analysis via the FindAllMarkers function from Seurat. Subsequently, we classified the cells into 7 distinct categories, based on the expression of canonical markers from relevant literature and expert knowledge. In addition, we implemented an unbiased automated annotation using the SingleR[45] package (V2.2.0). The reference for automated annotation was a log-normalized global cardiac atlas dataset from the human Heart Cell Atlas.

**Single-cell differential gene expression and pathway enrichment analysis**

We performed differential gene expression analysis (Seurat FindMarkers) within each unique cell type between two sets of conditions: E2 (ECDΔ/-) versus WT and E3 (ICDΔ/-) versus WT, using the WT data as the control. Differentially expressed genes were selected at a threshold of adjusted p-value < 0.05 and |log2FoldChange| > 0.25. Volcano plots highlighting the differentially expressed genes were generated using the EnhancedVolcano package (V1.18.0). Gene Set Enrichment Analysis (GSEA)[46] was performed using clusterProfiler[47](V4.9.0.2) to identify significantly enriched or suppressed pathways in E2 and E3 samples, compared to the WT samples. The following gene sets were utilized for the enrichment analysis, all of which were procured from Molecular Signature Database[48] via msigdbr (V7.5.1): KEGG, C2 (curated), and C5 (ontology) gene sets. Multiple testing was corrected using the Benjamini-Hochberg method; genes and pathways with an adjusted p-value < 0.05 were considered significant.

**Pseudotime trajectory analysis of single-cell data**

Pseudotime trajectory analysis was performed on the integrated dataset subset to the following clusters: Non-specific Proliferating, Fibroblast-like Proliferating, Cardiomyocyte, Smooth Muscle, and Fibroblast cells, utilizing the monocle3 package[49,50 51,52] (V1.3.1). The integrated Seurat dataset was converted into a Cell Data Set (CDS), a process which included the direct transfer of UMAP embeddings and cluster labels, as well as all relevant cell metadata. Strict quality control measures were implemented to ensure the accurate transfer of cell labels and metadata, including the verification of gene identifiers and cell barcode – metadata pairs. The analysis then followed the standard monocle3 procedure:

1. Potential cell trajectories were inferred from the reduced UMAP space using reversed graph embedding.
2. The earliest principal node, which serves as the origin for pseudotime calculations, was determined based on the set of potential principal nodes output and a selected “root” cluster (Non-specific Proliferating cells).
3. Cells were ordered in pseudotime along calculated trajectories.
4. 4 distinct lineages were programmatically identified via the choose_graph_segment function, defining a minimum spanning tree between the principal “root” node and a terminal “leaf” node.

5. Genes that were differentially expressed across a lineage were identified using the graph_test function. Markers with a Moran's I > 0.25 and q-value < 0.05 were identified to have a significant change in expression across the lineage.
6. Heatmaps highlighting genes with the most significant change in expression across the lineage, with cells ordered along the pseudotime lineage, were generated using ComplexHeatmap[53] (V2.16.0).

**Statistical analyses and reproducibility**

Statistical analyses were performed using R version 4.3.0, and all data used for this paper will be available in the GEO repository.

**Transfection**

Plasmid transfections were conducted using Lipofectamine 3000 transfection reagent.

**FDA-approved compound screening**

High throughput screening of FDA-approved compounds as conducted as previously described[54]. Briefly, a subset of the FDA-approved compound library (TargetMol) was stamped to a 384-well tissue plate (100nL per drug) using the CyBio Well vario (Analytik Jena). Compounds were plated to the final concentrations of 10 μM. Clonal NICD-HiBiT expressing BEAS-2B cells were seeded to final density of 3000 cells per well. After 18 h of treatment, culture media was removed and cells were processed for Nanoluciferase activity using Lytic HiBiT detection system (Promega), according to manufacturer's protocol. Cell viability was measured by CellTiter-Glo 2.0 (Promega). Signals were collected and quantified using a CLARIOstar plate reader (BMG Labtech).

**Immunoblotting**

Immunoblotting assays were conducted as previously described[54]. Briefly, cells were lysed in RIPA buffer supplemented with EDTA-free protease inhibitor cocktail. Lysate was sonicated and precipitated by centrifugation, and supernatant was normalized for total protein concentration and mixed with 6x Protein Sample Buffer. Processed lysate was then analyzed by SDS-PAGE Bolt gels (Invitrogen), and then electro-transferred to nitrocellulose. Blots were blocked using 20% milk in TBST for 1 h before primary antibody incubation overnight at 4 °C. Following washing, and secondary antibody incubation, blots were washed and visualized using the West Femto Maximum Sensitivity Substrate (Thermo Scientific) on a UVP ChemStudio Imager (AnalytikJena).
